# Environmental factors contribute to cancer therapy toxicity

**DOI:** 10.1101/2025.11.11.687867

**Authors:** Lindsey C. Mehl, Maria Sacconi Nuñez, Javier Marco Sanz, Anna Geraghty, Louisa Dal Cengio, Annika Kaval, Yishu Chen, Erin M. Gibson

**Affiliations:** Department of Psychiatry and Behavioral Sciences, Stanford University School of Medicine, Palo Alto, CA 94305, USA; Cancer Biology Graduate Program, Stanford University School of Medicine, Palo Alto, CA 94305, USA; Department of Pediatrics, Clinica Universidad de Navarra, Navarra, Spain; Howard Hughes Medical Institute, Stanford University, Stanford, CA 94305, USA; Department of Neurology and Neurological Sciences, Stanford University, Palo Alto, CA 94305, USA

## Abstract

Cancer therapy-related cognitive impairment (CRCI) affects numerous cancer patients, however there is substantial variability in its severity. Here, we investigated how environmental factors such as housing facility and time of treatment, known as chronotherapy, impact chemotoxicity in a pediatric murine model of methotrexate (MTX)-induced CRCI. We find that MTX consistently impacts body mass in a time-of-day specific manner across different housing facilities. Intriguingly, investigation of the gut microbiome acutely after MTX chronotherapy revealed substantial differences in microbial composition between animals housed in different facilities as well as decreased microbial diversity at different times of day. Furthermore, these differences in the gut microbiome after MTX chronotherapy coincide with differences in circulating inflammatory cytokine profiles between facilities and times of day. Chronically, housing facility continued to impact serum cytokine levels after MTX treatment, whereas the effect of time wanes. Nonetheless, we found that time drastically alters chronic white matter microglial gene expression in the central nervous system (CNS) after MTX treatment. Together, these findings demonstrate that both housing facility and time dictate response to MTX chronotherapy in the periphery and CNS. Collectively, this work elucidates putative factors that can regulate MTX-induced chemotoxicity.

## Introduction

Cancer therapy-related cognitive impairment (CRCI) is a neurological disorder associated with cancer treatment that manifests as deficits in learning, memory, attention, information processing, executive functioning, multitasking, speech/language skills, fine motor function, and mood disorders^1–4^. With more than 18.6 million current cancer survivors living in the United States and more than 22 million predicted cancer survivors by 2035^5^, more people than ever are at risk for developing CRCI. Despite recent advancements in understanding the etiology of CRCI^6^, there is considerable variability in how this debilitating condition presents across tumor types and treatment regimens. Studies of genetic vulnerability have identified variants in oxidative stress and metabolic genes that are associated with cognitive deficits after cancer treatment in humans and animal models^7–13^. Demographics like age^14^, sex, and hormone status have also been implicated in differential neurological dysfunction after cancer therapy^15–18^. However, comparatively less is known about how environmental factors impact the manifestation of chemotoxicity.

Novel research has begun to explore the relationship between CRCI and environmental factors such as diet^19,20^ and the gut microbiome^21,22^. The gut microbiome has been associated with modulating response to cancer treatment^23–26^ and mediating CRCI^27–30^. This work is galvanizing in light of emerging research suggesting that crosstalk between the gut and the brain can promote neuroinflammation and neurodegeneration^31–34^. Recent findings also indicate that, outside the context of cancer treatment, gut health in animal models is extensively affected by environmental factors such as housing conditions^35^ and the circadian system^36–38^. This work underscores the need to fully characterize how these environmental factors come together to regulate CRCI.

Circadian biology encompasses an expansive field of research studying how time of day impacts biological processes in both health^39^ and disease^40^. It is well characterized how the body responds to environmental cues such as light to regulate homeostatic processes like eating and cellular metabolism^41^. One corollary of circadian biology is chronotherapy, or timed administration of a treatment, which harnesses circadian oscillations in cellular homeostatic processes to maximize therapeutic potential. Cancer treatment chronotherapy has been studied extensively, but clinical application of chronotherapy has been slow and research has predominantly focused on therapeutic efficacy rather than toxicity^42–50^. Despite this, revolutionary work has found that time of day can control peripheral and CNS inflammation in response to cyclophosphamide and doxorubicin^51^. As such, there is a critical need to better understand how cancer chronotherapy affects toxicity to prioritize quality of life for cancer survivors worldwide.

Methotrexate (MTX) is an antimetabolite chemotherapy that has been used to treat cancer for decades, and continues to be a commonly utilized therapeutic agent, particularly for hematological malignancies^52,53^. Our previous work elucidated neuroinflammation as a driver of MTX-induced CRCI in a pediatric murine model. The study identified white matter-specific microglial reactivity as the lynchpin for a neuroinflammatory cascade resulting in long-term CRCI^54^. Early studies of MTX began to explore chronotherapy efficacy^55^ and toxicity^56,57^. Yet, these preclinical chemotoxicity studies were limited to adult male rodents and did not attempt to assess neurotoxicity associated with MTX chronotherapy. In light of this work, we hypothesized that environmental factors could modulate the severity of MTX-induced chemotoxicity, and we sought to investigate how factors such as time and housing facility may account for the heterogeneity in chemotoxicity with our juvenile MTX-induced CRCI model.

## Results

### Establishing a juvenile MTX chronotherapy murine model

Our previous work established a juvenile, MTX-induced CRCI murine model^54^. To determine how circadian biology may impact chemotoxicity, we adapted this dosing scheme to generate a juvenile MTX chronotherapy paradigm (Figure 1A). Our treatment regimen consisted of weekly injections of either 100 milligrams per kilogram (mg/kg) MTX or control phosphate buffered saline (PBS) administered intraperitoneally. Treatment was administered at either ZT0 (lights on, start of the murine rest phase) or ZT12 (12 hours later, lights off, start of the murine active phase) beginning at postnatal day 21 (P21). Mice received treatment once per week for 3 weeks, with the final injection on P35. Mice always received the same treatment at the same time of day for all 3 doses. To study the effect of animal housing facility, this injection paradigm was completed at two different low barrier animal housing facilities across Stanford campus. Both housing facilities were subject to the same standards of animal husbandry with the same food and water provided at both facilities.

**Figure 1.**
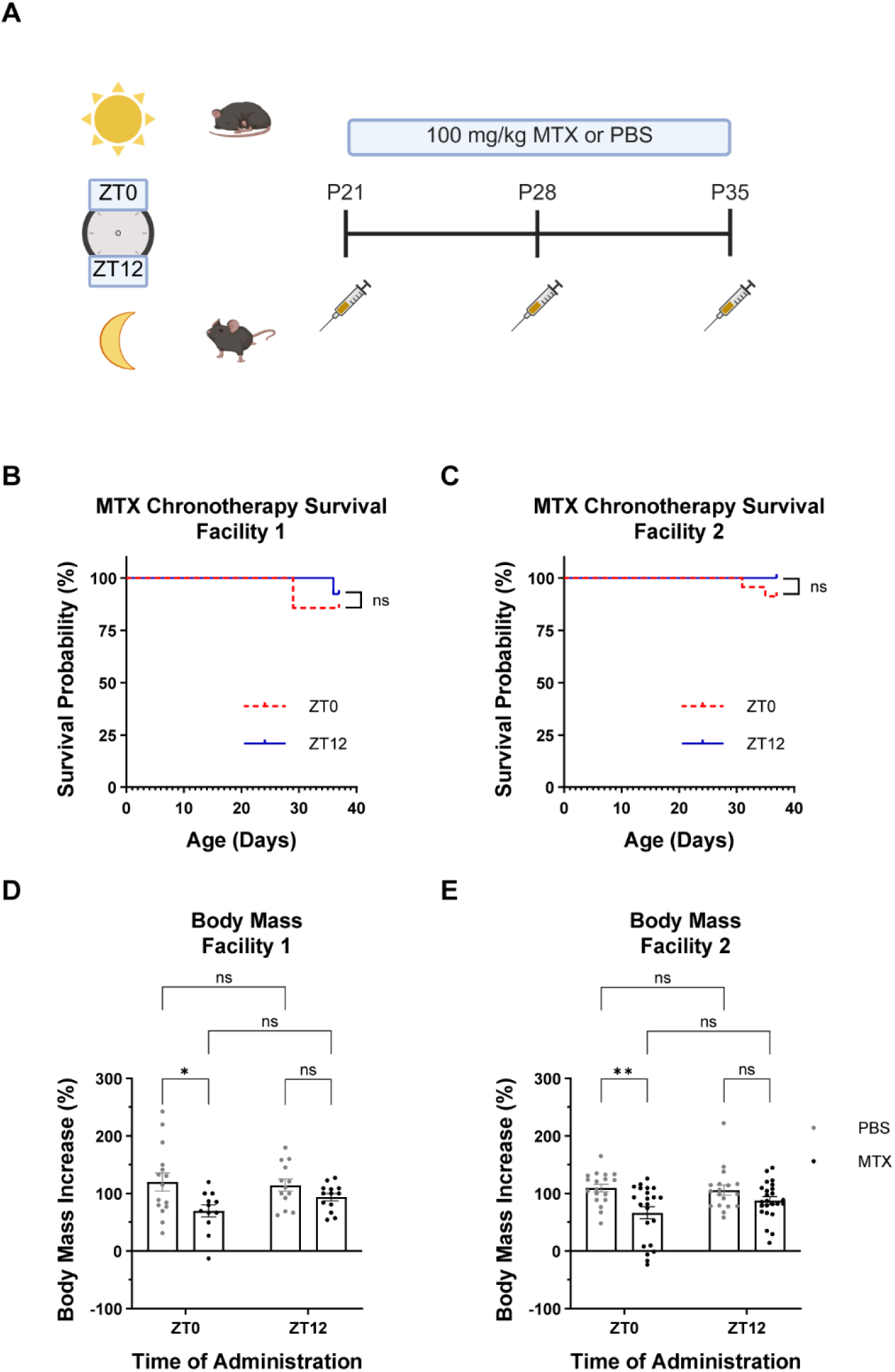
MTX timing does not impact survival but differentially impairs body mass increase. (A) Chronotherapy dosing paradigm for juvenile mice treated at either ZT0 (lights on) or ZT12 (lights off) with either 100 mg/kg MTX or PBS via intraperitoneal injection. Mice were treated once a week for 3 weeks beginning at postnatal day 21 (P21). This paradigm was completed across two different animal facilities at our home institution. Created in BioRender. Mehl, L. (2025) https://BioRender.com/beq2a3z. (B) Animals treated with MTX in Facility 1 show no difference in overall survival after completion of the chronotherapy treatment paradigm when they are treated at ZT0 versus ZT12 (ZT0, n = 14; ZT12, n = 13; p = 0.5718) by Log-rank (Mantel-Cox) test. (C) Animals treated with MTX in Facility 2 show no difference in overall survival after completion of the chronotherapy treatment paradigm when they are treated at ZT0 versus ZT12 (ZT0, n = 23; ZT12, n = 24; p = 0.1441) by Log-rank (Mantel-Cox) test. (D) Over the course of the chronotherapy treatment paradigm in Facility 1, animals treated with MTX at ZT0 did not gain as much mass compared to PBS treated animals at ZT0 (PBS, n = 15; MTX, n = 12; p = 0.0200) but there was no difference in the ability to gain mass between MTX versus PBS treated animals at ZT12 (PBS, n = 13; MTX, n = 13; p = 0.6105) by 2way ANOVA with Tukey’s multiple comparisons tests. (E) Over the course of the chronotherapy treatment paradigm in Facility 2, animals treated with MTX at ZT0 did not gain as much mass compared to PBS treated animals at ZT0 (PBS, n = 18; MTX, n = 22; p = 0.0033) but there was no difference in the ability to gain mass between MTX versus PBS treated animals at ZT12 (PBS, n = 18; MTX, n = 24; p = 0.4108) by 2way ANOVA with Tukey’s multiple comparisons tests. Column graph data shown as mean ± SEM. n.s. p > 0.05; *p < 0.05; **p < 0.01; ***p < 0.001; ****p < 0.0001

To assess the viability of our cross-facility, MTX chronotherapy model, we first characterized basic survival metrics across the different housing facilities and treatment administration times. Work in a preclinical rat model suggests that MTX chronotherapy may affect treatment tolerance^57^, and MTX maintenance chronotherapy in human children with pediatric leukemia has been shown to have worse disease-free survival outcomes when administered in the morning compared to the evening^55^. Though our animal model does not include a tumor component so that we can specifically dissect the effects of chronotherapy treatment, we sought to rigorously interrogate whether MTX chronotherapy had adverse effects on survival. Fortunately, MTX treatment at ZT0 versus ZT12 does not affect overall survival outcomes in Facility 1 (Figure 1B) or Facility 2 (Figure 1C). As expected, mice treated with control PBS at ZT0 versus ZT12 show no differences in overall survival over the course of the chronotherapy treatment paradigm in either facility (Figure S1A-B). There were also no adverse effects on survival when comparing MTX versus PBS treatment at any combination of administration time or facility (Figure S1C-F). These data suggest that MTX administration at both ZT0 and ZT12 is well tolerated and has translational potential.

While MTX chronotherapy did not affect overall survival, we anecdotally observed that MTX animals treated at ZT0 appeared consistently smaller than their PBS control treated counterparts. We found that, in both Facility 1 and Facility 2, ZT0 MTX treated animals failed to gain as much body mass as their time-matched PBS treated littermates (Figure 1D-E) across the treatment paradigm. During this phase of life, healthy juvenile mice typically gain mass rapidly, and the failure to do so suggests that time of day may play a key role in peripheral toxicity associated with our MTX chronotherapy model. This finding is also consistent with previous work in male adult rodents, which found that MTX chronotherapy can affect body mass^56^.

### Housing facility impacts the gut microbiome after MTX chronotherapy

Considering the consistent body mass deficits experienced by ZT0 MTX-treated animals and that gastrointestinal toxicity is the most common adverse side effect of MTX cancer treatment in humans^58,59^, we next sought to understand how MTX chronotherapy may be affecting the gut microbiome of animals undergoing our cross-facility MTX chronotherapy paradigm. We used shotgun metagenomic sequencing to elucidate the gut microbial composition of animals treated with MTX or PBS at ZT0 and ZT12 across both housing facilities at the same institution. At 48-hours after the final injection, fecal samples were collected at the same time as injection. Fecal samples were then submitted to TransnetYX (Cordova, TN) for library preparation and sequencing (Figure S2A, Methods). Shotgun sequencing of fecal samples allowed us to compare genus-level taxonomic differences and microbial diversity between facilities after MTX chronotherapy.

To determine the state of the gut microbiome after MTX chronotherapy, we first assessed the average relative abundance of the top 20 genera in fecal samples collected from MTX-treated mice housed in Facility 1 (Figure 2A) versus Facility 2 (Figure 2B). We used the MaAsLin2^60^ R package to identify genera that were significantly affected by the housing facility and determine the facility effect size for each individual genus. MaAsLin2^60^ relies on a general linear model, and we treated both Facility and Time as independent fixed effects for this analysis. Strikingly, taking into account both ZT0 and ZT12, we identified 7 genera (*Sangeribacter, Bacteroides, Lactobacillus, Bifidobacterium, Alistipes, Paramuribaculum,* and *Kingevirus*) that were all significantly affected by facility, despite the facilities being located at the same institution and subject to the same husbandry standards (Q < 0.05, Figure 2C-D, Table S1). Five genera, including *Bacteroides, Lactobacillus, Alistipes, Paramuribaculum, and Kingevirus,* had a higher relative abundance in Facility 1, whereas *Sangeribacter* and *Bifidobacterium* had a higher relative abundance in Facility 2. Notably, both the *Sangeribacter*^61^ and *Bifidobacterium*^62^ genera have been associated with a healthy gut microbiome. We next asked whether facility impacts the diversity of microbes after MTX chronotherapy regardless of time, given that microbial diversity has been linked to cancer treatment response^23,25,26^. We used the OneCodex platform to compute each sample’s alpha diversity, which takes into account the richness and evenness of microbe distribution within samples. Intriguingly, when assessing alpha diversity with the Simpson index, we observe decreased alpha diversity in Facility 2 compared to Facility 1. However, when assessing the Shannon Index of alpha diversity, we see a similar trend, but it is not significant (Figure 2E). This could perhaps be attributed to differences between these diversity metrics in sensitivity to the presence of rare microbes versus changes in dominant microbes.

**Figure 2.**
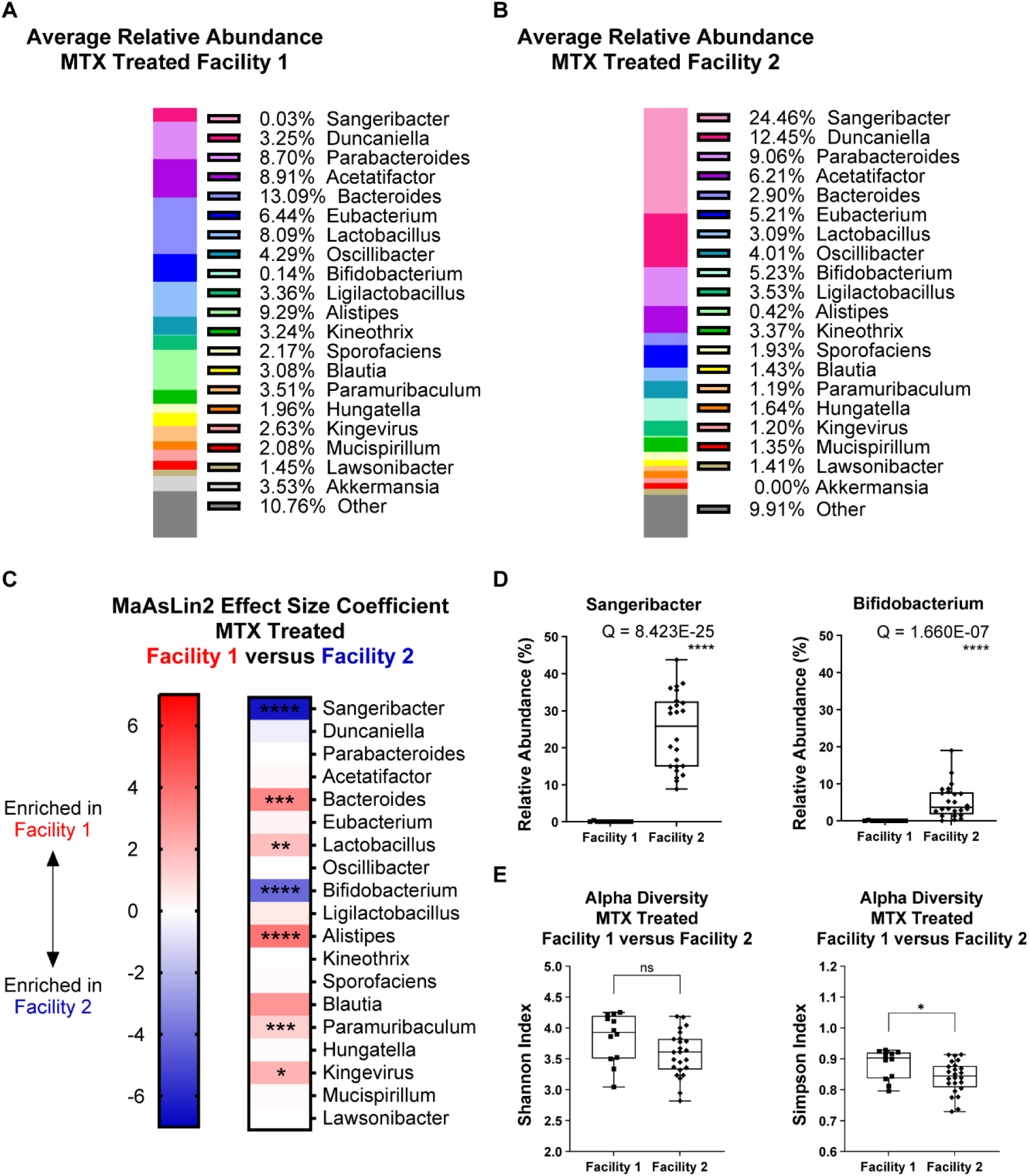
Housing facility alters microbiome composition after MTX chronotherapy. Animals were treated with MTX chronotherapy until P35 in either Facility 1 or Facility 2, and fecal pellets were collected at ZT0 and ZT12 on P37 for shotgun metagenomic microbiome sequencing. Sequencing reads were assigned to specific microbial genera and the relative abundance of microbes in each sample was computed using the OneCodex platform. The difference between microbial composition in Facility 1 versus Facility 2 after MTX treatment was assessed with the MaAsLin2 R package^60^ (See methods). (A) Average relative abundance of the top 20 microbial genera for MTX treated Facility 1 fecal samples (n = 12). (B) Average relative abundance of the top 20 microbial genera for MTX treated Facility 2 fecal samples (n = 24). (C) MaAsLin2 effect size coefficient was plotted as a heat map for each genus, with positive effect sizes (red) indicating enrichment in Facility 1 and negative effect sizes (blue) indicating enrichment in Facility 2 (Facility 1, n = 12; Facility 2, n = 24). Stars denote Q values of genera that were significantly affected by facility. A list of effect sizes and Q values for each genus can be found in Table S1. (D) Representative box and whisker plots of the relative abundance for genera significantly impacted by facility after MTX treatment as assessed by MaAsLin2 (Facility 1, n = 12; Facility 2, n = 24; Sangeribacter, Q = 8.423E-25; Bifidobacterium, Q = 1.660E-07). (E) The alpha diversity (both Shannon and Simpson indices) of each fecal sample was computed with the OneCodex platform for samples from Facility 1 versus Facility 2 after MTX treatment. There was not a significant difference in alpha diversity when looking at the Shannon Index (Facility 1, n = 12; Facility 2, n = 24; p = 0.0600) by unpaired two-tailed t test. However, there was a significant difference between Facility 1 and Facility 2 when alpha diversity was measured with the Simpson Index (Facility 1, n = 12; Facility 2, n = 24; p = 0.0181) by Mann-Whitney test. n.s. p > 0.05; *p < 0.05; **p < 0.01; ***p < 0.001; ****p < 0.0001 n.s. Q > 0.05; *Q < 0.05; **Q < 0.01; ***Q < 0.001; ****Q < 0.0001

### Time influences the gut microbiome after MTX chronotherapy

Our MaAsLin2^60^ analysis also explored the gut composition in MTX chronotherapy samples collected at ZT0 (Figure 3A) versus ZT12 (Figure 3B) to understand whether time of day also impacts the microbiome after MTX chronotherapy. Taking into account both facilities, we found that time significantly affected the relative abundance of 12 of the top 20 microbial genera (Q < 0.05, Figure 3C-D, Table S2). At ZT0 we observed a greater relative abundance of *Acetatifactor, Eubacterium, Oscillibacter, Kineothrix, Sporofaciens, Hungatella, Mucispirillum,* and *Lawsonibacter.* In contrast, at ZT12 we identified a greater relative abundance of *Bacteroides, Lactobacillus, Alistipes, and Kingevirus*. We investigated if alpha diversity also differs throughout the day in both facilities. We found that samples collected at ZT0 had significantly higher alpha diversity scores than ZT12 samples when assessed with both the Shannon and Simpson indices (Figure 3E). This suggests that the alpha diversity of samples does indeed change during the day, with greater diversity being observed at ZT0 (end of active phase) than at ZT12 (end of rest phase), which may impact the chemotoxicity associated with MTX treatment. These findings additionally highlight how critical collection time is when assessing chemotoxicity.

**Figure 3.**
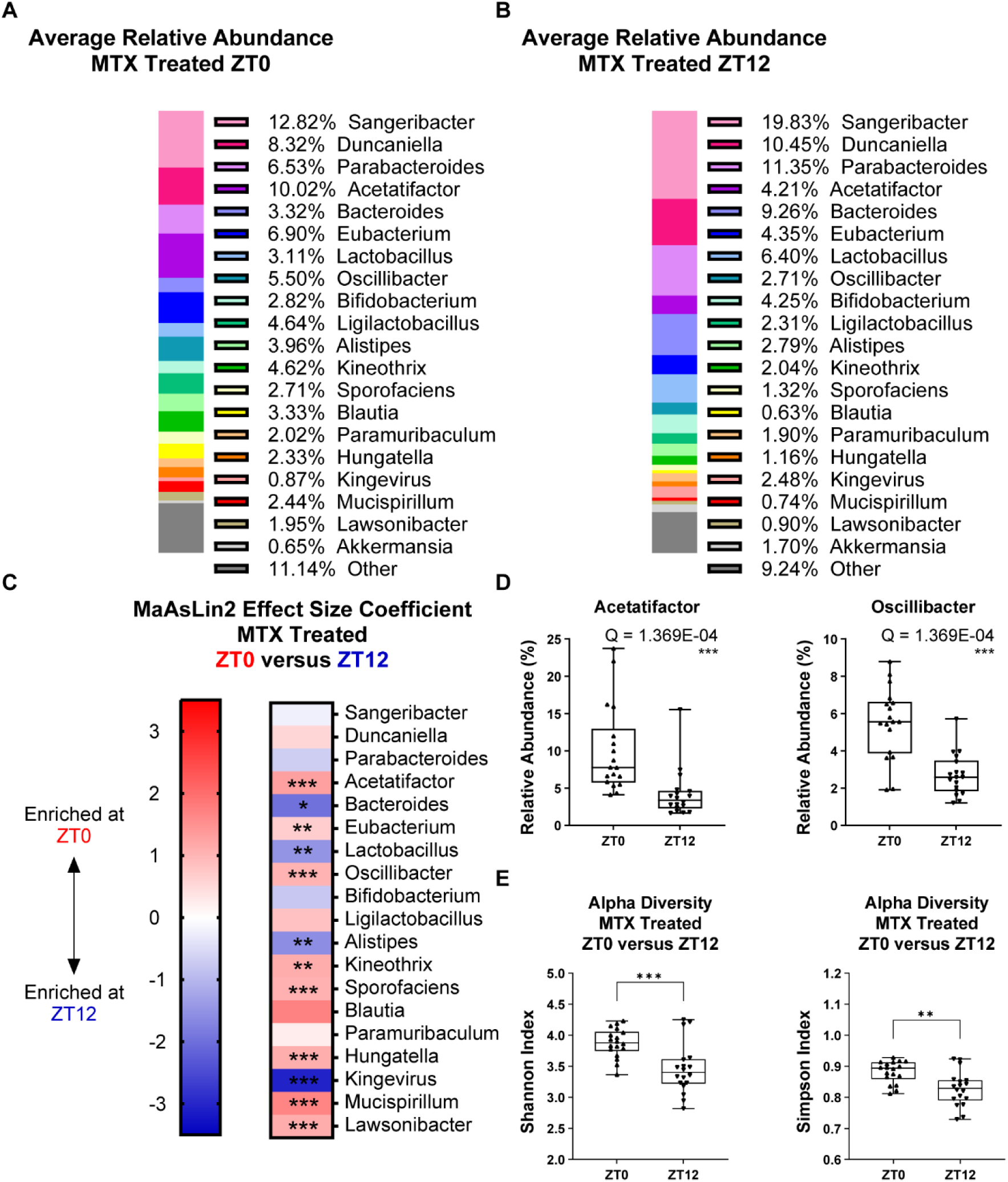
Time of day controls microbiome composition after MTX chronotherapy. Animals were treated with MTX chronotherapy until P35 in Facility 1 and Facility 2, and fecal pellets were collected at ZT0 or ZT12 on P37 for shotgun metagenomic microbiome sequencing. Sequencing reads were assigned to specific microbial genera and the relative abundance of microbes in each sample was computed using the OneCodex platform. The difference between microbial composition at ZT0 versus ZT12 after MTX chronotherapy was assessed with the MaAsLin2 R package^60^ (See methods). (A) Average relative abundance of the top 20 microbial genera for MTX treated ZT0 fecal samples (n = 18). (B) Average relative abundance of the top 20 microbial genera for MTX treated ZT12 fecal samples (n = 18). (C) MaAsLin2 effect size coefficient was plotted as a heat map for each genus, with positive effect sizes (red) indicating enrichment at ZT0 and negative effect sizes (blue) indicating enrichment at ZT12 (ZT0, n = 18; ZT12, n = 18). Stars denote Q values of genera that were significantly affected by time. A list of effect sizes and Q values for each genus can be found in Table S2. (D) Representative box and whisker plots of the relative abundance for genera significantly impacted by time after MTX treatment as assessed by MaAsLin2 (ZT0, n = 18; ZT12, n = 18; Acetatifactor, Q = 1.369E-04; Oscillibacter, Q = 1.369E-04). (E) The alpha diversity (both Shannon and Simpson indices) of each fecal sample was computed with the OneCodex Platform for samples at ZT0 versus ZT12 after MTX treatment. Alpha diversity was consistently higher at ZT0 versus ZT12 after MTX treatment when looking at both the Shannon Index (ZT0, n = 18; ZT12, n = 18; p = 0.0007) and the Simpson Index (ZT0, n = 18; ZT12, n = 18; p = 0.0013) by unpaired two-tailed t tests. n.s. p > 0.05; *p < 0.05; **p < 0.01; ***p < 0.001; ****p < 0.0001 n.s. Q > 0.05; *Q < 0.05; **Q < 0.01; ***Q < 0.001; ****Q < 0.0001

### Both housing facility and time modulate the gut microbiome at baseline

To determine how housing facility and time impact the gut microbiome at baseline, we also sequenced and analyzed microbial composition in PBS treated animals across facilities and administration times. As with the MTX chronotherapy samples, we first used OneCodex to compute the average relative abundance of genera in the gut microbiome after control PBS treatment in Facility 1 (Figure S2B) and Facility 2 (Figure S2C), taking into account both ZT0 and ZT12. We then used MaAsLin2 again (with housing facility and time as independent, fixed effects) to assess the effect size and significance of housing facility on specific genera. Even at baseline, with only control PBS injections, the relative abundance of 6 genera (*Bacteroides, Sangeribacter, Lactobacillus, Bifidobacterium, Alistipes,* and *Paramuribaculum*) was significantly affected by facility (Figure S2D-E, Table S3). Genera including *Bacteroides, Lactobacillus, Alistipes,* and *Paramuribaculum* had higher relative abundance in Facility 1 while *Sangeribacter* and *Bifidobacterium* had greater relative abundance in Facility 2. Despite the significant effect of facility on the relative abundance of 6 genera across times, there was no difference in overall alpha diversity in Facility 1 versus Facility 2 (Figure S2F).

We next computed the average relative abundance of PBS treated samples at ZT0 (Figure S3A) and ZT12 (Figure S3B) taking both facilities into account. MaAsLin2 analysis of ZT0 versus ZT12 samples yielded that collection time significantly impacted the relative abundance of 11 different genera including *Bacteroides, Acetatifactor, Eubacterium, Parabacteroides, Kineothrix, Oscillibacter, Alistipes, Sporofaciens, Hungatella, Lawsonibacter,* and *Dorea* (Figure S3C-D, Table S4). When we computed alpha diversity at ZT0 versus ZT12 in both facilities with OneCodex, we again found that samples collected at ZT0 showed significantly higher diversity when assessed with both the Shannon and Simpson indices (Figure S2E). Together, these results indicate that facility and time are both key regulators of the microbial composition and diversity.

### Both housing facility and time determine serum cytokines acutely after MTX chronotherapy

Emerging work has begun to elucidate the critical role of the gut-brain axis in modulating proper neural function and neurodegeneration, particularly via peripheral inflammation^31–33^. Given that our MTX chronotherapy CRCI model demonstrates stark differences in microbial composition across facilities and at different times of day, we next asked about the state of circulating serum cytokines after MTX chronotherapy. As with our fecal collection paradigm (Figure S2A), we began by treating mice weekly with 100 mg/kg MTX at either ZT0 or ZT12 starting on P21. 48-hours after the final injection on P37, at either ZT0 or ZT12, we collected blood from the saphenous vein and isolated the serum. We then performed a fluorescence-based assay to measure the levels of 48 different murine cytokines (Figure S4A, Methods). Median fluorescence intensity (MFI) readings were processed into a batch corrected measures of cytokine level (dpMFI) with a R utility^63^ developed at the Stanford Human Immune Monitoring Center to correct for batch effects and nonspecific binding artifacts prior to all downstream analyses.

To elucidate how specific cytokine MFIs vary after MTX chronotherapy, we completed a cytokine profiling analysis^64^ on all MTX treated samples collected at P37. We analyzed these data with a centered polynomial restricted maximum likelihood model (REML), a mixed effects linear model. The results of the family level tests for each fixed effect and interaction can be found in Table S5. As expected, cytokine level (dpMFI) significantly varied depending on the cytokine (p < 0.0001, Table S5), which validates that different cytokines were present in the serum at different levels. Cytokine dpMFI also significantly varied by Facility alone (p = 0.0013, Table S5), regardless of cytokine identity, suggesting a potent effect of facility on peripheral inflammation. This led us to examine the interaction between Facility and Cytokine.

Our Cytokine profile analysis revealed a significant interaction between Cytokine and Facility (p < 0.0001, Table S5), and we computed the least squares mean estimates and standard error for the difference between Facility 1 and Facility 2 as well as the p value for each individual cytokine. Taking into account both ZT0 and ZT12, this analysis identified 6 cytokines whose levels were significantly affected by interaction with facility: Leptin, CCL4, IL2, IL6, CCL5, and CXCL10 (Figure 4A, Table S6). Intriguingly, all 6 cytokines of interest were upregulated in Facility 1 relative to Facility 2. Leptin is particularly intriguing given its role in appetite since ZT0 MTX treatment impaired body mass increase across both facilities (Figure 1D-E). Furthermore, leptin has been shown to have a role in inflammation^65^ and can be influenced by the microbiome^66^. Additionally, IL2, IL6, CCL4, and CCL5 are all generally proinflammatory, perhaps suggesting a more robust peripheral immune response in Facility 1 versus Facility 2 acutely after MTX treatment. Notably, CXCL10 has also been identified as a cytokine of interest in the context of chimeric antigen receptor T (CART) cell therapy-induced CRCI, which has a similar neuroinflammatory mechanism to MTX-induced CRCI^67^.

**Figure 4.**
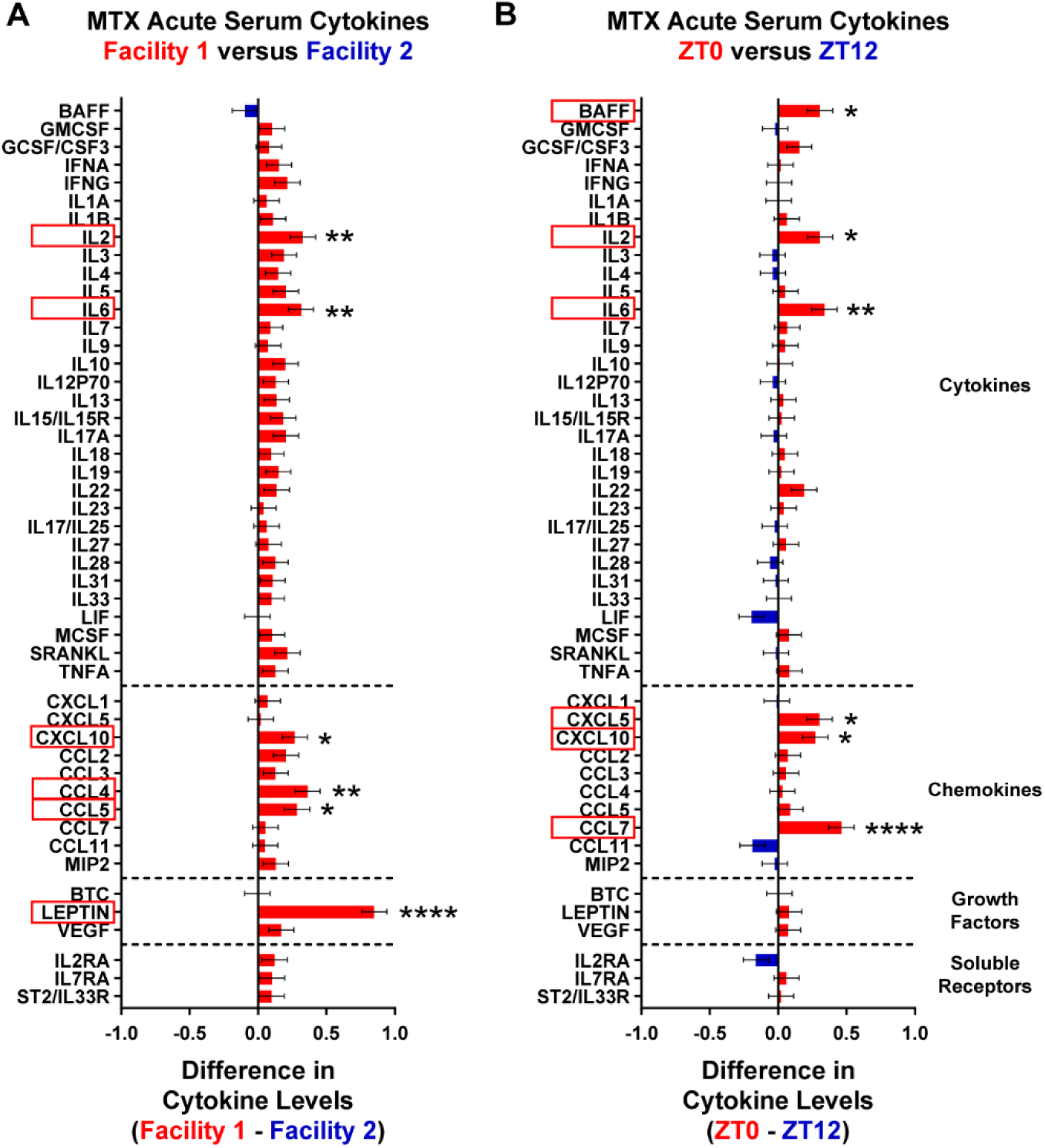
Housing facility and time dictate acute peripheral inflammation after MTX chronotherapy. Animals were treated with MTX chronotherapy in Facility 1 or Facility 2, and blood was collected from the saphenous vein at either ZT0 or ZT12 on P37 at the same time as injection to assess peripheral inflammation. P37 MTX treated serum cytokine levels were analyzed via cytokine profiling analysis with an REML mixed effects model with Facility, Time, and Cytokine as fixed effects (See methods). (A) For MTX-treated samples collected at P37, a significant interaction between Facility and Cytokine was identified by cytokine profile analysis (Facility 1, n = 23; Facility 2, n = 23; p = <0.0001; Table S5). P values for interactions between Facility and individual cytokines were FDR-adjusted, and cytokines with Q values less than 0.05 were considered significantly affected by interaction with Facility. Figure displays the difference in least squares mean estimates of Facility 1 versus Facility 2 (Facility 1, n = 23; Facility 2, n = 23; Leptin, Q = 1.87E-17; CCL4, Q = 0.00221; IL2, Q = 0.00660; IL6, Q = 0.00814; CCL5, Q = 0.0196; CXCL10, Q = 0.0286). Values greater than zero indicate higher expression in Facility 1 (red), and values less than zero indicate higher expression in Facility 2 (blue). Data shown as ± SE. All cytokine results reported in Table S6. (B) For MTX-treated samples collected at P37, a significant interaction between Time and Cytokine was identified by cytokine profile analysis (ZT0, n = 23; ZT12, n = 23; p = <0.0001; Table S5). P values for interactions between Time and individual cytokines were FDR-adjusted, and cytokines with Q values less than 0.05 were considered significantly affected by interaction with Time. Figure displays the difference in least squares mean estimates of ZT0 versus ZT12 (ZT0, n = 23; ZT12, n = 23; CCL7, Q = 3.17E-05; IL6, Q = 0.00613; BAFF, Q = 0.0109; CXCL5, Q = 0.0109; IL2, Q = 0.0109; CXCL10, Q = 0.0269). Values greater than zero indicate higher expression at ZT0 (red), and values less than zero indicate higher expression at ZT12 (blue). Data shown as ± SE. All cytokine results reported in Table S7.

The cytokine profile analysis of acute serum cytokines after MTX chronotherapy in both facilities also yielded a significant interaction between Cytokine and Time (p < 0.0001, Table S5), and we again computed the least squares mean estimates and standard error for the difference between ZT0 and ZT12 as well as the p value for each individual cytokine interaction with Time. We identified another 6 cytokines that were significant (Q < 0.05) and upregulated at ZT0, including CCL7, IL6, BAFF, CXCL5, IL2, and CXCL10 (Figure 4B, Table S7). Like CXCL10, CCL7 was also implicated in CART-induced CRCI^67^. The other cytokines are also associated with inflammatory immune cell recruitment, which is particularly intriguing as peripheral immune activation can in turn promote neuroinflammation^68–71^. These findings are also concordant with previous studies in adult male rodents that identified fluctuations in leukocyte numbers resulting from MTX chronotherapy^56,57^. Collectively, the cytokine profile analysis indicates that, like the gut microbiome, peripheral inflammation can be impacted by both housing facility and time acutely after MTX chronotherapy.

### Only housing facility regulates serum cytokine levels chronically after MTX chronotherapy

We repeated the same above-mentioned cytokine profiling analysis with MTX samples collected at P65. As anticipated, cytokine dpMFI values significantly varied as a result of Cytokine identity (p < 0.0001, Table S5). Similar to P37, the P65 family level test for the interaction between Facility and Cytokine was significant (p < 0.0001, Table S5). At this chronic timepoint 1-month post-treatment, we computed the least squares mean estimate and standard error for the difference between Facility 1 and Facility 2 as well as the p value for each individual cytokine. Taking both times into account, we only identified 3 individual cytokines that had a significant interaction with Facility (Q < 0.05). Nonetheless, these 3 cytokines (Leptin, CCL4, and CXCL10), were all more highly expressed in Facility 1 than Facility 2 (Figure 5A, Table S8). This result posits a persistent peripheral immune response that is stronger in Facility 1 compared to Facility 2. Strikingly, when taking both facilities into account, the family level test for the interaction of Time and Cytokine was not significant (p = 0.119, Table S5). However, the difference in least squares means estimates between ZT0 and ZT12 show a trend toward increased expression at ZT0 (Figure 5B, Table S9). Furthermore, the 3-way interaction of Facility, Time, and Cytokine was significant (p < 0.0001, Table S5), suggesting that circulating cytokine levels vary specifically due to the combination of Cytokine, Facility, and Time. The specific cytokines significantly affected by the 3-way interaction (Q < 0.05) included IL2, Leptin, CCL4, CXCL10, IL22, and SRANKL, confirming that both Facility and Time are crucial regulators of the systemic immune response after MTX chronotherapy (Table S10).

**Figure 5.**
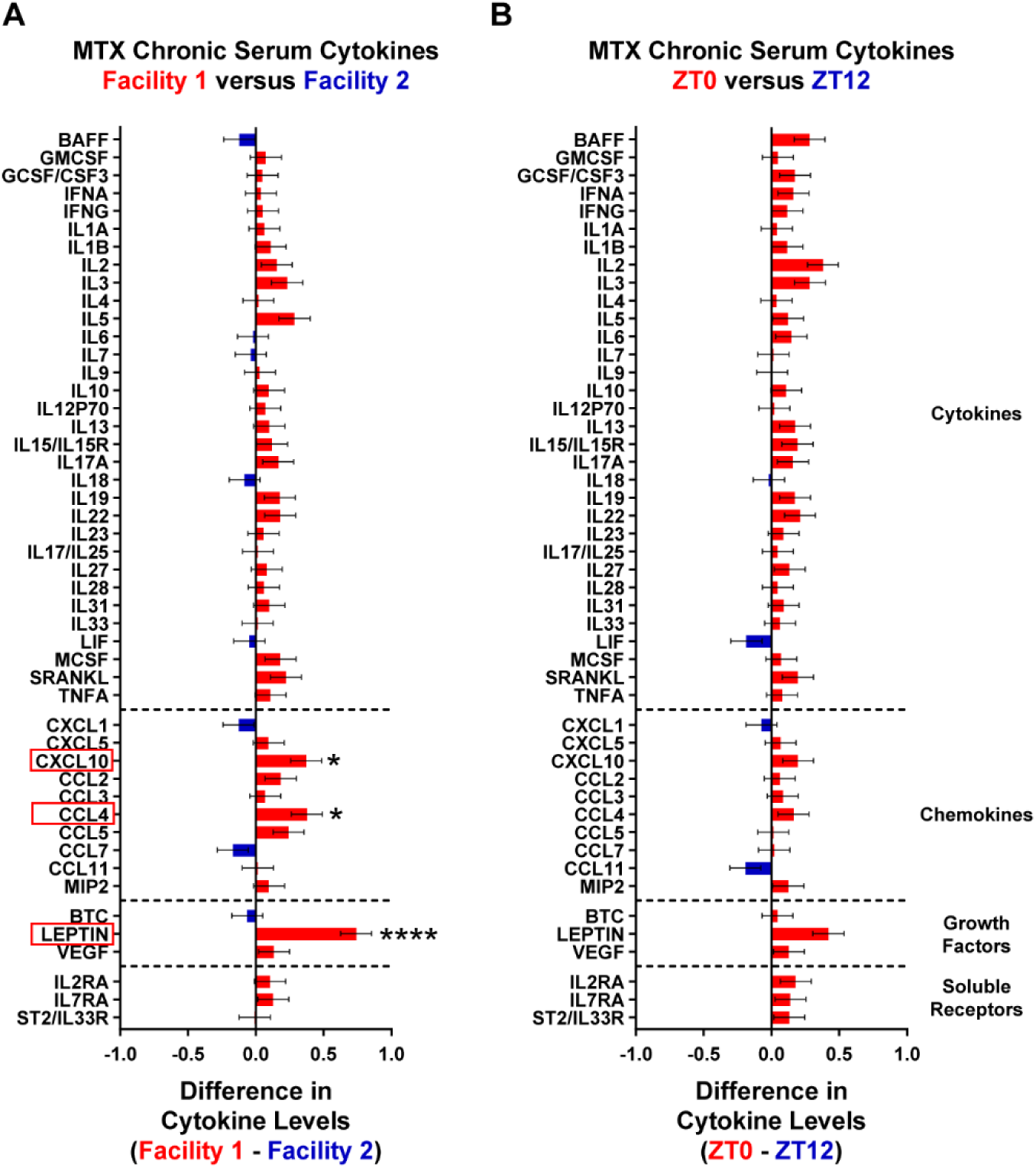
Housing facility has modest effect on chronic peripheral inflammation after MTX chronotherapy. Animals were treated with MTX chronotherapy in Facility 1 or Facility 2, and blood was collected from the saphenous vein at either ZT0 or ZT12 on P65 at the same time as injection to assess peripheral inflammation. P65 MTX treated serum cytokine levels were analyzed via cytokine profiling analysis with an REML mixed effects model with Facility, Time, and Cytokine as fixed effects (See methods). (A) For MTX-treated samples collected at P65, a significant interaction between Facility and Cytokine was identified by the cytokine profile analysis (Facility 1, n = 19; Facility 2, n = 18; p = <0.0001; Table S5). P values for interactions between Facility and individual cytokines were FDR-adjusted, and cytokines with Q values less than 0.05 were considered significantly affected by interaction with Facility. Figure displays the difference in least squares mean estimates of Facility 1 versus Facility 2 (Facility 1, n = 19; Facility 2, n = 18; Leptin, Q = 1.13E-08; CCL4, Q = 0.0196; CXCL10, Q = 0.0196). Values greater than zero indicate higher expression in Facility 1 (red) and values less than zero indicate higher expression in Facility 2 (blue). Data shown as ± SE. All cytokine results reported in Table S8. (B) For MTX-treated samples collected at P65, the interaction between Time and Cytokine was not significant (ZT0, n = 19; ZT12, n = 18; p = 0.119; Table S5). As such, FDR-adjusted p values (Q values) were not used to determine significant interactions between Time and individual cytokines. Figure displays the difference in least squares means estimates of ZT0 versus ZT12. Values greater than zero indicate higher expression at ZT0 (red), and values less than zero indicate higher expression at ZT12 (blue). Data shown as ± SE. All cytokine results reported in Table S9. n.s. Q > 0.05; *Q < 0.05; **Q < 0.01; ***Q < 0.001; ****Q < 0.0001

### Both housing facility and time impact acute and chronic inflammation at baseline

To determine the effects of housing and time on circulating cytokine levels at baseline, we repeated the cytokine profiling analysis, but this time with blood samples collected at P37 after PBS chronotherapy (Figure S4A). Again, cytokine dpMFI levels significantly varied depending on the cytokine (p < 0.0001, Table S5). At this acute timepoint, cytokine dpMFI levels also varied as a result of Facility alone (p < 0.0001, Table S5), regardless of cytokine identity, which drove us to investigate the interaction of Facility and Cytokine at baseline. Indeed, when considering both ZT0 and ZT12, the interaction of Cytokine and Facility was also significant (p < 0.0001, Table S5), and we computed the least squares mean estimate and standard error for the difference between Facility 1 and Facility 2 as well as the p value for each individual cytokine interaction with Facility. Astoundingly, at baseline we identified 21 cytokines that were significantly affected by interaction with Facility (Q < 0.05), which was nearly half the cytokines in the panel (Figure S4B, Table S11). The majority of these cytokines (19 out of 21) had higher expression in Facility 1 than Facility 2, suggesting that the animals in Facility 1 are at baseline in a more primed, proinflammatory state than animals in Facility 2.

When taking into account both facilities, we also observed a significant interaction between Time and Cytokine (p < 0.0001, Table S5) and further analysis of the individual cytokines and time identified 4 cytokines of significance: IL2, CCL11, IL6, and CCL2 (Figure S4C, Table S12). These cytokines are particularly interesting, with both CCL11 and CCL2 being implicated in CART-induced CRCI^67^, and CCL11 also being associated with microglial reactivity and cognitive impairment^72,73^. Together the results suggest that time of day also influences baseline immune activity. Furthermore, the family level test of the 3-way interaction of Facility, Time, and Cytokine was also significant (p < 0.0001, Table S5), suggesting that cytokine levels vary depending on the cytokine identity and the combination of Facility and Time. Individual cytokines that varied with both Facility and Time (Q < 0.05) were Leptin, IL2, IL6, IL3, CCL2, CCL11, CCL7, IFNA, IFNG, IL5, IL17A, IL22, CCL5, CCL4, GMCSF, IL10, and IL1A, which includes many cytokines broadly considered proinflammatory (Table S13).

We repeated the cytokine profiling analysis a final time on PBS-treated samples collected at P65 in both facilities at both times to understand factors that affect peripheral inflammation at baseline at a later developmental age (Figure S5A, Table S14). As with previous analyses, the family level test for Cytokine was significant (p < 0.0001, Table S5). The interaction of Cytokine and Facility was also significant (p < 0.0001, Table S5) and further analysis in SAS identified just 2 cytokines, Leptin and IL3, that significantly varied (Q <0.05) due to their interaction with Facility. Both were upregulated in Facility 1 compared to Facility 2 (Figure S5B). The interaction of Cytokine and Time was also significant (p < 0.0039, Table S5), though the assessment of the interactions between time and individual cytokines yielded only IL2 as significantly affected by the interaction with Time (Figure S5C, Table S15). Finally, the interaction of Facility and Time (independent of specific cytokine) was unexpectedly significant (p = 0.021, Table S5). Further studies may shed more light on the complex relationship between serum cytokines, housing facility, and time of day throughout aging.

### Time modulates long-term microglial gene expression after MTX chronotherapy

Due to the differential peripheral inflammation we observed after MTX chronotherapy (Figures 4-5) and recent work highlighting the relationship between the microbiome, peripheral inflammation, neuroinflammation, and CRCI^28^, we next decided to examine the CNS in our model of CRCI induced by MTX chronotherapy. We focused on white matter microglia because our previous work identified white matter microglia as gatekeepers of a brain region-specific neuroinflammatory cascade caused by MTX^54^. To accomplish this, we conducted bulk RNA-sequencing of microglia from white matter murine brain tissue after MTX chronotherapy treatment in Facility 2. At 1-month post-treatment, the corpus callosum was microdissected, the white matter was dissociated, and microglia were enriched via magnetic activated cell sorting (MACS, Figure S6A). Total RNA was isolated from these microglial samples and submitted to Novogene Co., LTD for library preparation and bulk RNA-sequencing. We first assessed the quality and similarity of biological replicates with a Pearson correlation between all samples. All samples had R^2^ values were greater than 0.96, indicating high similarity (Figure S6B). We repeated the Pearson Correlation amongst biological groups and observed R^2^ values greater than 0.98 (Figure S6C). Finally, we completed a Principal Component Analysis (PCA) to examine the distribution of biological replicates from each biological group prior to completing downstream analyses (Figure S6D).

After these quality controls, we investigated the relative gene expression of MTX treatment at ZT0 versus ZT12 and identified nearly 500 differentially expressed genes (DEGs, Q < 0.05, Figure 6A). Of these DEGs, there were 121 upregulated and 356 downregulated at ZT0 compared to ZT12. Notably, among our most significant hits were core circadian clock genes including *Arntl* (the gene encoding BMAL1, the primary circadian timekeeper), Period family genes (*Per2* and *Per3*), and Rev-Erb family genes (*Nr1d1* and *Nr1d2*). These DEGs bolstered our confidence that baseline time of day biology was intact and not disrupted by the microglial isolation process. We performed Gene Ontology (GO) pathway enrichment analysis on the DEGs between MTX at ZT0 versus ZT12. For the pathways that were significant (Q < 0.05), we identified those with the highest gene ratio and plotted these in a scatter plot (Figure 6B). As expected, rhythmic processes were among our top pathways as this pathway includes classic circadian genes (i.e. *Dbp, Per3, Nr1d2, Nr1d1, Per2,* and *Arntl*). Intriguingly, the significant pathways with the highest gene ratios also included those relating to the extracellular matrix, cell migration, and lipid localization, which suggests that time of day may differentially impact microglial movement after MTX chronotherapy. To get a sense of how time impacts gene expression at baseline, we also assessed differential microglial gene expression of PBS treated animals 1-month after treatment at ZT0 versus ZT12 (Figure S6E). DEGs again included key circadian genes, but the total number of DEGs, was only about 36 genes in total. As pathway analysis ideally includes hundreds to thousands of genes, we did not perform further GO analysis. Taken together, these results indicate that time substantially regulates the gene expression of white matter microglia after MTX chronotherapy.

**Figure 6.**
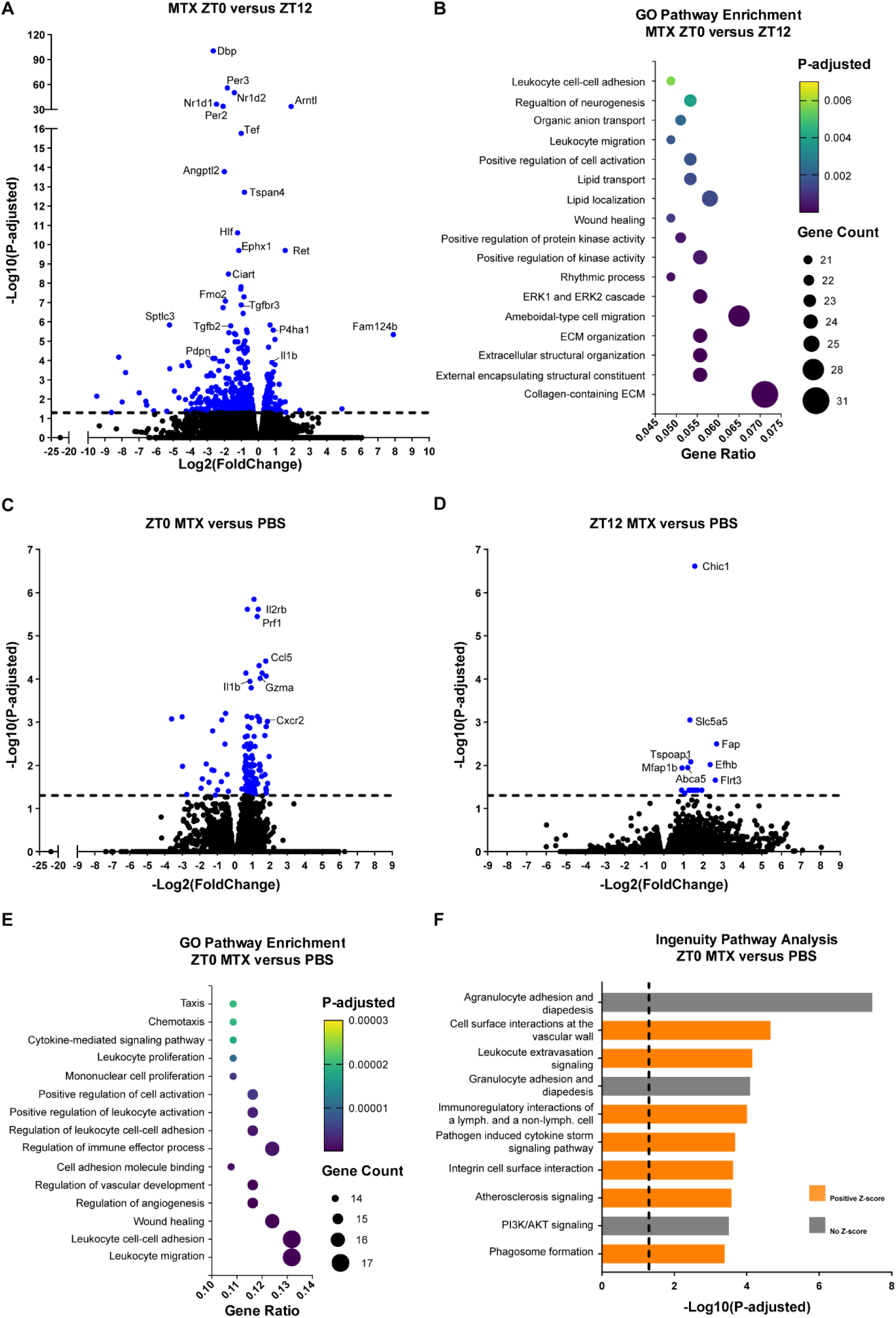
MTX chronotherapy differentially affects microglial inflammatory gene expression. Bulk RNA-sequencing (RNA-seq) was performed at 1-month post-treatment on MACS enriched microglia from the corpus callosum of mice that underwent the chronotherapy treatment paradigm in Facility 2. (A) Volcano plot showing differentially expressed genes in MTX ZT0 versus ZT12 samples. Genes with P-adjusted (Q value) less than 0.05 are shown in dark blue (n = 3 biological replicates per biological group). (B) Gene ontology (GO) pathways enriched in differentially expressed genes from MTX ZT0 versus ZT12 samples. Plotted pathways have P-adjusted (Q value) less than 0.05 and were pathways with the highest gene ratios. (C) Volcano plot showing differentially expressed genes in ZT0 MTX versus PBS samples. Genes with P-adjusted (Q value) less than 0.05 are shown in dark blue (n = 3 biological replicates per biological group). (D) Volcano plot showing differentially expressed genes in ZT12 MTX versus PBS samples. Genes with P-adjusted (Q value) less than 0.05 are shown in dark blue (n = 3 biological replicates per biological group). (E) Gene ontology (GO) pathways enriched in differentially expressed genes in ZT0 MTX versus PBS samples. Plotted pathways have P-adjusted (Q value) less than 0.05 and were pathways with the highest gene ratios. Multiple pathways are related to pro-inflammatory response and cellular migration after ZT0 MTX treatment relative to ZT0 PBS control. (F) Qiagen Ingenuity Pathway Analysis showing pathways significantly enriched in differentially expressed genes in ZT0 MTX versus PBS samples. Z-score indicates directionality of pathway enrichment with a positive z-score indicating that the pathway is upregulated in ZT0 MTX samples relative to ZT0 PBS samples. No Z-score indicates the directionality of the pathway enrichment is undetermined. Significantly upregulated pathways are related to a pro-inflammatory response and cellular migration after ZT0 MTX treatment relative to ZT0 PBS controls.

### Chronic microglial reactivity ensues after ZT0 MTX chronotherapy

To further elucidate how MTX chronotherapy impacts microglial gene expression, compared MTX microglial gene expression to time-matched PBS controls. We assessed white matter microglial gene expression at either ZT0 (Figure 6C) or ZT12 (Figure 6D). Strikingly, at ZT0, we identified 134 DEGs (115 upregulated, 19 downregulated) by MTX treatment. However, at ZT12, we only identified 16 DEGs total. We completed GO pathway enrichment analysis on ZT0 DEGs and plotted significant (Q < 0.05) pathways with the highest gene ratio. Excitingly, many of these pathways revealed differences in immune cell, specifically leukocyte, migration and activation (Figure 6E). GO pathway enrichment identifies pathways of interest but does not take into account the direction of DEG fold change. As such, we further completed Qiagen Ingenuity Pathway Analysis (IPA) to determine whether these immune activation pathways were upregulated or downregulated by MTX treatment. IPA again identified a number of significant pathways associated with leukocyte migration and proinflammatory immune cell crosstalk and further determined that these pathways are upregulated in MTX treated samples compared to PBS controls at ZT0 (Figure 6F). In summary, bulk RNA-sequencing analyses of white matter microglia illuminated that ZT0 MTX chronotherapy specifically induced chronic microglial reactivity compared to time-matched controls.

## Discussion

Factors regulating variability in CRCI severity are incompletely understood. Here, we investigated the effects of housing facility and chronotherapy in a juvenile mouse model of MTX-induced CRCI to explore the extent to which environmental components can impact chemotoxicity. We identified both housing facility and time as major modulators of the gut microbiome, which coincided with altered systemic inflammation acutely after MTX chronotherapy. Chronically, housing facility persistently impacted circulating cytokines, and time dictated differential microglial gene expression following MTX chronotherapy. Ultimately, we pinpointed that ZT0 MTX chronotherapy specifically induced chronic microglial dysregulation relative to control PBS treatment that is not observed at ZT12. Ultimately, this work comprehensively characterizes the chemotoxicity associated with MTX chronotherapy and reveals environmental factors that may account for the heterogeneity associated with the severity of MTX chemotoxicity. Moving forward, it is critical to consider these factors when translating CRCI preclinical models into strategies for alleviating CRCI in cancer survivors.

Our finding that housing facility has profound effects on gut health corroborates work at other institutions^35^ and contributes to a larger conversation about irreproducibility in scientific research^74–76^. This knowledge is pivotal given that further research has demonstrated that the gut microbiome can alter cancer treatment response^23–26^. Two genera, *Sangeribacter* and *Bifidobacterium*, had higher relative abundance in Facility 2 than Facility 1 (Figure 2). Notably, *Sangeribacter* and *Bifidobacterium* have been associated with a healthy gut^61^ and colitis resolution^62^, respectively. Furthermore, *Sangeribacter* belongs to the Muribaculaceae family, which has been shown to help restore gastrointestinal health after MTX treatment in rats^77^. This raises the intriguing possibility that animals housed in Facility 2 may be better equipped to weather cancer therapy treatment.

Indeed, one species of *Bifidobacterium* has been associated with ameliorating liver and gastrointestinal toxicity caused by irinotecan chemotherapy via downregulating proinflammatory cytokines^78^. *Bifidobacterium* treatment has also been shown to decrease circulating cytokine levels and attenuate autoimmunity after anti-cancer checkpoint blockade^79^. This is concordant with our findings that circulating cytokine levels are higher in Facility 1 than Facility 2 acutely (Figure 4) and chronically (Figure 5) after MTX. Some of the cytokines elevated in Facility 1, such as CXCL10, have also been implicated in CART-induced CRCI^67^. Others, such as leptin, have been tied to the microbiome as well^66^. Strikingly, we observed numerous cytokines acutely affected by facility after PBS treatment alone, even more than after MTX treatment. It is possible that proinflammatory effects triggered by high dose MTX, in contrast to the anti-inflammatory effects of low dose MTX^80^, may partially obfuscate the differences between facilities. The differential peripheral inflammation between facilities is notable for our MTX chronotherapy CRCI model because prior work has established that the immune system can act as a conduit along the gut-brain axis^31–33^. Independent of the microbiome differences between facilities, systemic inflammation has been linked to the onset of neuroinflammation in a number of disease contexts^68–71^. Collectively, this body of work supports collaborative multicenter studies across which findings are consistently validated.

Our findings also identify time as a potent modulator of microbial composition and circulating serum cytokine levels after MTX and PBS treatment. Our results that time of day impacts the microbiome and its diversity aligns with other studies that have enshrined circadian dynamics regulating microbial composition and metabolism^36,37,81^. In turn, microbial depletion can reciprocally disrupt host circadian synchrony as well^82^. Unfortunately, collecting fecal and blood samples at the same time as treatment complicates disambiguating “collection time” versus “treatment time” effects after MTX chronotherapy. However, our comparison of control PBS at ZT0 versus ZT12 (Figure S3-S5) gives us a clear idea of the effect of collection time for both microbiome composition and circulating cytokines. Both the MTX and PBS chronotherapy comparisons between ZT0 and ZT12 demonstrate that time of day significantly impacts both the gut microbiome and systemic inflammation in the short-term after treatment.

Analysis of serum samples at our long-term timepoint after MTX chronotherapy indicated that expression of some circulating cytokines continued to be significantly affected by facility (Figure 5) but not by time. The waning impact of interaction with time perhaps suggests a decrease in peripheral inflammation by P65, which would not be surprising given the relatively short half-life of MTX^83^. Intriguingly though, in the CNS, there are still numerous genes that were differentially expressed between microglia from animals treated with MTX chronotherapy at ZT0 versus ZT12 at this one-month post-treatment (Figure 6A). These genes convey differences in microglial mobility, extracellular matrix navigation, and lipid transport across the day (Figure 6B). This may suggest that these microglia are distinct after MTX chronotherapy because this was not apparent when analyzing microglial gene expression in ZT0 versus ZT12 PBS treated animals. To get at this question further, we examined the gene expression of MTX versus PBS treated animals at ZT0 and again at ZT12 and found differential upregulation of proinflammatory pathways only at ZT0 (Figure 6C-F). Thus, future studies will aim to identify mechanistic pathways and other neural cell types involved in perturbing chronic microglial gene expression after MTX chronotherapy and will investigate facility effects on microglial state changes.

### Limitations

Though microdissection of the corpus callosum prior to microglial isolation is essential to capturing white matter-specific microglial gene expression via bulk RNA-sequencing, it limits the amount of total microglial RNA that can be collected. MACS isolation via CD11b beads was the optimal microglia isolation strategy given the time sensitive nature of the protocol and the availability of equipment at specific times of day. While all samples were enriched for microglia via MACS before total RNA extraction, there is likely RNA contribution from other CNS cell types (and border-associated macrophages) present in the samples. A critical caveat of this work is that the B6 wild-type sires of experimental mice for each facility came from different vendors. It is true that animal vendor can have an effect of experimental outcomes^84–86^. However, all mice used in our experiments were from the F1 generation of B6 males crossed to CD1 females that were purchased from the same vendor. Therefore, experimental mice lived in their respective facilities for their entire lives. Furthermore, others have found that fecal microbial composition in low-barrier facilities like where our experimental mice are housed shift within weeks of arrival from the vendor. The composition after this initial shift then remains relatively stable in the subsequent F1 and F2 generations^35^. Given that our breeding animals are given weeks to acclimate to our facility before experiments proceed and that we only conduct experiments with the F1 generation of animals that live their entire lives in the same facility, we believe microbiome and inflammation differences between facilities are a result of the facility conditions rather than vendor differences.

## Materials

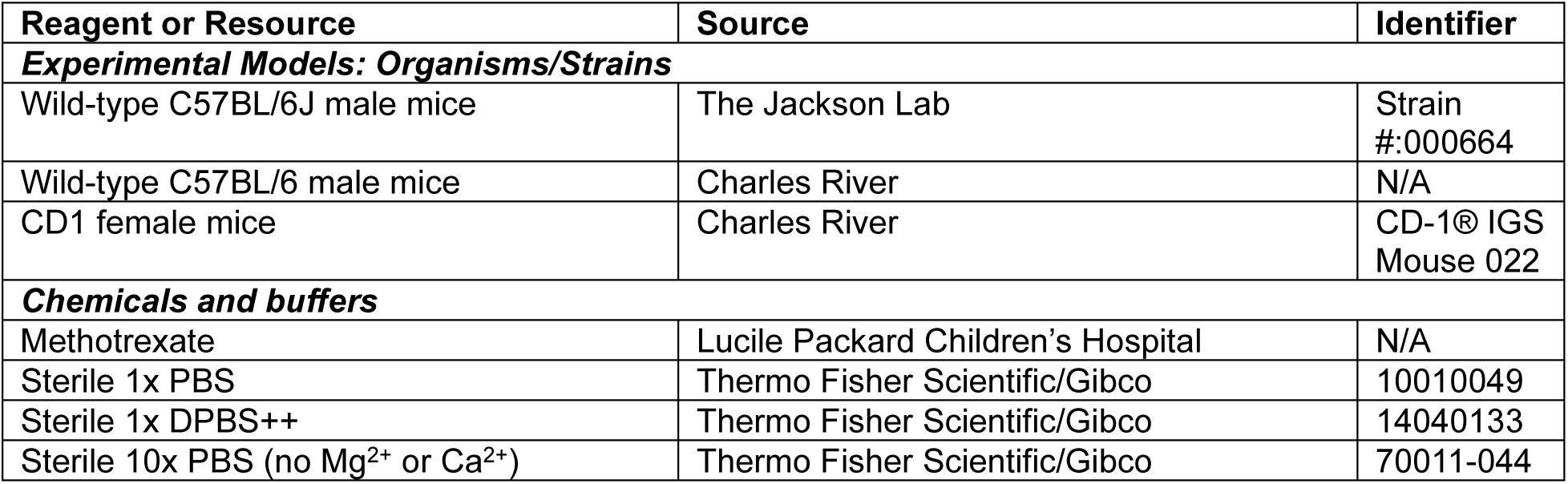
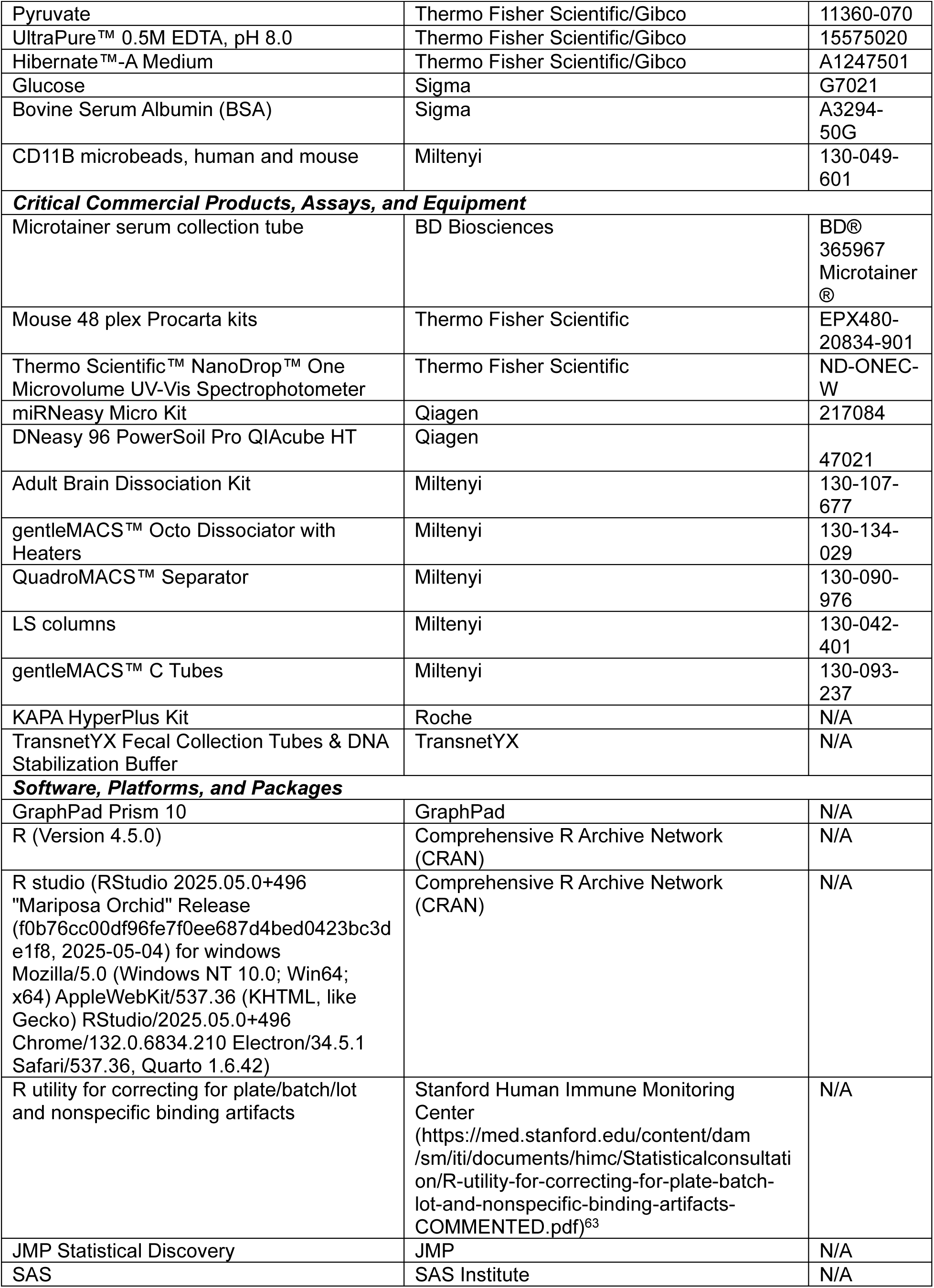
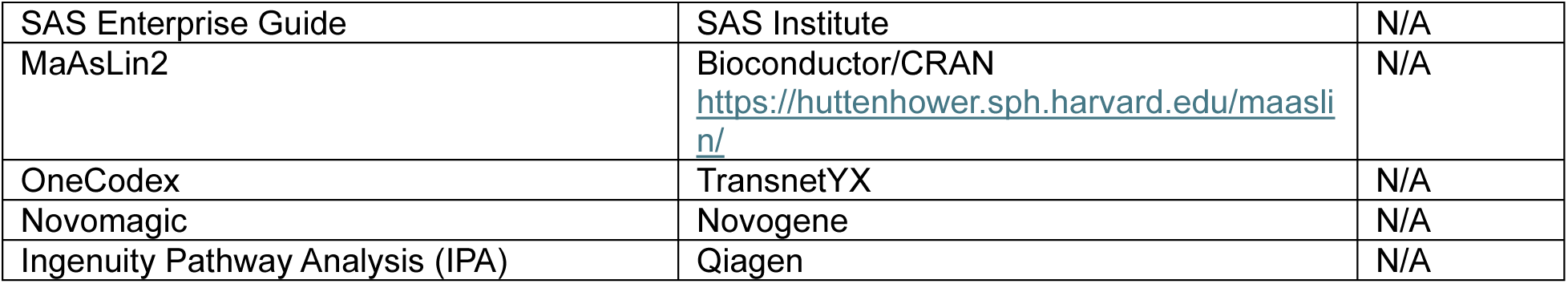

## Methods

### Mouse maintenance

In Facility 1, wild-type C57BL/6 male mice (Charles River) were bred with CD1 female mice (Charles River) for all experiments. In Facility 2, wild-type C57BL/6J male mice (The Jackson Lab) were bred with CD1 female mice (Charles River) for all experiments. Each animal studied was of the F1 generation of this cross. Each animal studied was of the F1 generation of this cross. All animals were housed in a 12-hour light, 12-hour dark cycle with *ad libitum* access to both food and water. Both sexes were used for all studies and housed with 2-5 animals per cage when possible. Mice were housed in the same cage with other animals of the same sex receiving the same treatment at the same time of day. Mice received only their assigned treatment at their assigned time of day and were not otherwise manipulated. All procedures were performed according to guidelines established by the Stanford University Institutional Care and Use Committee (IACUC). Mice were housed in two different housing facilities on Stanford campus, referred to as Facilities 1 and 2, and received the same husbandry practices. Animals in both facilities were subject to the same protocols described above.

### Chronotherapy treatment paradigm

C57BL/6J:CD1 mice of both sexes were given a weekly intraperitoneal (IP) injection of phosphate buffered saline (PBS) or 100 mg/kg methotrexate (MTX) dissolved in sterile PBS at P21, P28, and P35 at either ZT0 or ZT12. All mice were massed the day prior to injection to determine dose volume of a 10 mg/mL MTX working stock solution. All procedures were performed according to guidelines established by our Stanford Administrative Panel of Laboratory Animal Care (APLAC) protocol. MTX stock (25 mg/mL) was obtained from the Lucile Packard Children’s Hospital at Stanford University and diluted to 10 mg/mL with sterile PBS to create a working stock prior to injection. Animals were sacrificed at either P37 (acute timepoint) or one-month post-treatment (∼P62-65, chronic timepoint) depending on the experimental endpoint. For all experiments mice were sacrificed at the same time of day as injection.

### Body mass analysis

Change in body mass was assessed across the injection paradigm with the initial body mass being recorded on P20, prior to the first injection, and the final body mass being recorded on P34 prior to final injection. Percent increase in body mass from initial measure was calculated and differences between biological groups were assessed via 2way ANOVA with Tukey’s multiple comparisons tests.

### Shallow shotgun metagenomic sequencing of murine fecal samples

Microbiome sequencing was performed in collaboration with TransnetYX (Cordova, TN, USA). Prior to fecal collection, plastic containers for fecal collection were autoclaved to ensure sterility. For all experiments, feces were collected at the same time of day as chronotherapy injections. At P37 at ZT0 or ZT12, after mice of both sexes were exposed to the same chronotherapy paradigm at P21-35 as described above, each animal was allowed to defecate into a sterile container. Using a sterile applicator, each fecal sample was transferred to a barcoded collection tube containing room temperature stable, DNA stabilization buffer provided by TransnetYX. Approximately 2-5 fecal pellets were collected for each animal. The barcoded TransnetYX collection tube with DNA stabilization buffer was then closed tightly and shaken to ensure the fecal pellets were fully covered by stabilization buffer. Between each animal, the collection area and personal protective equipment were cleaned with 70% ethanol. A new sterile plastic container was used for each animal. All samples were stored protected from light at room temperature until all samples were collected and submitted to TransnetYX for DNA extraction, library preparation, sequencing, and read assignment. Once TransnetYX received the fecal samples, they extracted inhibitor-free, high molecular weight genomic DNA with the Qiagen DNeasy 96 PowerSoil Pro QIAcube HT extraction kit and protocol. After performing DNA extraction and DNA quality control (QC) assessment, TransnetYX prepared sequencing libraries from genomic DNA with the KAPA HyperPlus library preparation protocol. TransnetYX then added unique dual indexed (UDI) adapters to ensure proper read assignment. After the standard library QC, the libraries were shotgun sequenced on an Illumina NextSeq 2000 at a sequencing depth of 2 million 2×150 base pair end reads. The DNA sequencing files were uploaded to the OneCodex analysis software and analyzed against the OneCodex Database of microbial genomes. The OneCodex Database consists of ∼148,000 complete microbial genomes, including ∼71,000 distinct bacterial genomes, ∼72,000 viral genomes, and thousands of archaeal and eukaryotic genomes assembled from both publicly and privately available sources. The OneCodex Database routinely undergoes automated and manual curation steps to remove low quality or mislabeled records. TransnetYX used reference mouse genomes to remove host DNA reads. TransnetYX then filtered the microbial classification results through several statistical post-processing steps to eliminate false positive results caused by contamination or sequencing artifacts. Relative abundance and alpha diversity (Shannon and Simpson indices) were obtained and visualized from the OneCodex analysis software before being exported for further plotting in GraphPad Prism 10 and statistical analysis in R. Alpha diversity scores were tested for normality with the Shapiro-Wilk and Kolmogorov-Smirnov tests. If alpha diversity scores passed normality tests, a parametric unpaired two-tailed t test was used to determine significance. If alpha diversity scores failed normality tests, a nonparametric Mann-Whitney test was used to determine significance. Significant differences between the microbial composition at ZT0 versus ZT12 and Facility 1 versus Facility 2 were determined with the MaAsLin2 R package (Mallick et al. 2021^60^), which relies on a general linear model. Time of day and facility were assigned as independent fixed effects (ZT12 and Facility 2 were set as references for Facility and Time, respectively) for the relative abundance of the 20 most prevalent microbial genera, and the MaAsLin2 package was run with default parameters. MaAsLin2 computes the P adjusted (Q value) of individual genera significantly affected by Facility or Time with the Benjamini-Hochberg method. Q values less than 0.05 indicated that a particular microbial genus was significantly affected by either time or facility. The MaAsLin2 coefficient (effect size) was plotted as a heatmap. MTX treated samples and PBS treated samples were analyzed separately.

### Serum collection and cytokine profiling statistical analyses

For all experiments, blood was collected at the same time of day as chronotherapy injections. At either P37 or 1-month post-treatment (∼P65), after mice of both sexes were exposed to the same chronotherapy paradigm at P21-35 as described above, serum was collected to assess the level of circulating cytokines. Cages were placed on a heating pad. Each animal was then properly restrained, and fur was shaved as necessary for a clear view of the saphenous vein. Alcohol wipes were used to clean the skin, and Vaseline was applied to the puncture site. A 25G needle was used to draw blood at a right angle to the vein, and blood was collected in a microtainer serum collection tube (BD Biosciences, BD® 365967 Microtainer®). The tubes were incubated at room temperature for at least 30 minutes but no more than 2 hours. The serum samples were placed in a microcentrifuge and spun down for 10 minutes at maximum speed at room temperature. The supernatant (serum) was removed and transferred to a sterile eppendorf before being stored at −80C. Serum circulating cytokines were assessed with the Luminex – Thermo-Fisher/Life technologies Mouse 48 plex Kits. This assay was performed by the Human Immune Monitoring Center at Stanford University. Mouse 48 plex Procarta kits (EPX480-20834-901) were purchased from Thermo-Fisher/Life Technologies, Santa Clara, California, USA, and used according to the manufacturer’s recommendations with modifications as described. Briefly, beads were added to a 96 well plate and washed in a BioTek ELx405 washer. Samples were added to the plate containing the mixed antibody-linked beads and incubated overnight at 4C with shaking. Cold (4C) and room temperature incubation steps were performed on an orbital shaker at 500-600 revolutions per minute (rpm). Following the overnight incubation, plates were washed in a BioTek ELx405 washer and biotinylated detection antibody was added for 60 minutes at room temperature with shaking. The plate was washed as described and streptavidin-PE was added for 30 minutes at room temperature. The plate was washed as above and reading buffer was added to the wells. Each sample was measured in duplicate. Plates were read using a Luminex 200 or a FM3D FlexMap instrument with a lower bound of 50 beads per sample per cytokine. Custom Assay Chex control beads were purchased from Radix BioSolutions (Georgetown, Texas) and added to all wells. Raw median fluorescence intensities (MFIs) for each cytokine in each sample were then taken by the core. Upon completion of all samples, the MFI values (from any combination of treatment (PBS or MTX), facility (1 or 2), age (P37 or P65), and time (ZT0 or ZT12)) were corrected for batch and nonspecific binding artifacts and log transformed with an R Utility developed at the Stanford HIMC core (Maecker et al., 2020^63^). The detrended preprocessed MFI (dpMFI) values then underwent proteomics profiling analysis (Coden et al. 2025^64^). P37 (acute) and P65 (chronic) samples were analyzed separately. MTX treated and PBS treated samples were also analyzed separately. Thus, we completed 4 analyses total: P37 MTX treated samples, P37 PBS treated samples, P65 MTX treated samples, and P65 PBS treated samples.

Cytokine profiling analyses in JMP and SAS used a centered polynomial restricted maximum likelihood (REML) mixed effects model. Our dependent Y variable was the batched corrected MFI (dpMFI) values for each of the 48 cytokines in each serum sample. We included Cytokine (called SP for soluble protein in the SAS code), Facility, and Time as independent fixed effects. We also included all 2-way interactions of these fixed effect terms (Cytokine and Facility, Cytokine and Time, Facility and Time) and the 3-way interaction of Cytokine, Facility, and Time as fixed effects. Finally, the model included specimen (individual mouse serum sample) nested within Facility and Time as a random effect such that each specimen served as its own control as described by Coden and colleagues (this method controls for unspecified variables such as cage barcode as well as inter-specimen variation)^64^. Fixed effect interactions passed a family level test (p < 0.05) before proceeding to test for significant interactions between a fixed effect and specific individual cytokines. SAS code and input data for each analysis are available upon request. If a 2-way or 3-way interaction including Cytokine passed the family level test, the p values for interactions between fixed effects and individual cytokines were FDR-adjusted with the Benjamini-Hochberg method (Q value) in JMP. Individual cytokines with Q values less than 0.05 were considered significantly affected by interaction with facility, time, or both facility and time. The difference in least squares mean estimate between facilities or times was plotted as a bar graph.

### Bulk RNA-Sequencing of corpus callosum microglia

For all experiments, brains were collected at the same time of day as chronotherapy injections. At 1-month post-treatment at ZT0 and ZT12, after all mice were exposed to the same chronotherapy paradigm at P21-35 as described above, mice of both sexes were sacrificed and their brains were extracted and placed in Hibernate A (ThermoFisher Scientific/Gibco, A1247501) until all brains for the timepoint had been extracted. When possible, separate tools were used for MTX versus PBS samples. If not possible, tools were cleaned with 70% ethanol between brain extractions for MTX and PBS-treated animals. The corpus callosum (CC) was microdissected and returned to Hibernate A to keep hydrated. All CCs from the same treatment condition were combined in a single eppendorf for a single biological replicate. Each biological replicate represents a separate litter of mice. CCs were spun down in a 4C microcentrifuge at 200g for 2 minutes. The supernatant was discarded, and the CCs were resuspended in approximately 1mL cold DPBS++ (ThermoFisher Scientific/Gibco, 14040133) to rinse. CCs were spun down in a 4C microcentrifuge at 200g for 2 minutes once more. The DPBS++ rinse was discarded, and CCs were transferred to gentleMACS™ C tubes for enzymatic dissociation with the Miltenyi Adult Brain Dissociation Kit (Miltenyi, 130-107-677) and gentleMACS™ Octo Dissociator with Heaters (Miltenyi, 130-134-029). CCs were mechanically dissociated with scalpels and scissors in EZ1 in the C tube and the remainder of brain dissociation proceeded according to manufacturer instructions (DPBS wash solution contained DPBS++, pyruvate (ThermoFisher Scientific/Gibco, 11360-070), and glucose (Sigma, G7021) as described in manufacturer instructions). Once dissociated, samples were enriched for microglia with magnetic activated cell sorting (MACS). MACS buffer (10x PBS no calcium or magnesium (ThermoFisher Scientific/Gibco, 70011-044), sterile water, EDTA (ThermoFisher Scientific/Gibco,15575020), BSA (Sigma, A3294-50G)) was kept on ice and positive selection for microglia was performed with CD11B microbeads (Miltenyi, 130-049-601) and LS columns (Miltenyi, 130-042-401). All microcentrifugation steps (10,000 rpm for 30s) were completed at 4C. The cells were incubated with CD11B microbeads on ice for 15 minutes. The LS columns were equilibrated with 2-3 washes of cold MACS buffer. After incubation, the cells were resuspended in cold MACS buffer, microcentrifuged to pellet, washed with cold MACS buffer once more, resuspended in cold MACS buffer a final time, and applied to the LS columns. The columns were washed 3 more times with cold MACS buffer before removal from the QuadroMACS™ Separator (Miltenyi, 130-090-976) magnet and elution in cold MACS buffer with the LS column plunger. The microglia were pelleted via centrifugation (300G, 10 minutes at 4C), resuspended in 500uL of Qiazol Lysis Reagent, and flash frozen on dry ice before storage at −80C. RNA was extracted with the Qiagen miRNeasy Micro Kit (Qiagen, 217084) using manufacturer’s instructions, and RNA quantity, quality, and purity were assessed with a Thermo Scientific™ NanoDrop™ One Microvolume UV-Vis Spectrophotometer. Total RNA was shipped on dry ice to Novogene Co., LTD for library preparation and bulk RNA sequencing. Prior to library preparation, Novogene assessed RNA quantity, integrity, and purity on an Agilent 5400 Fragment Analyzer system. Each sample included in library preparation passed QC with more than 100ng of RNA starting material and a RIN greater than 7. Messenger RNA was purified from total RNA using poly-T oligo-attached magnetic beads by Novogene. After fragmentation, first strand cDNA was synthesized using random hexamer primers, followed by the second strand cDNA synthesis using either dTTP for non-strand specific library or dUTP for strand specific library. The sequencing library was checked with Qubit and real-time PCR for quantification and then checked with bioanalyzer for size distribution. After library QC was completed, libraries were pooled based on the effective concentration and targeted data amount (9G of raw data per sample) and then sequenced on the Illumina NovaSeq™ X Plus Platform, which generated 150bp paired end reads. Novogene then processed raw fastq reads through the fastp software to remove reads containing adapters, reads containing ploy-N and low-quality reads. The Q20, Q30 and GC content were ascertained from the cleaned reads. All the downstream analyses were generated with these high-quality cleaned reads. Hisat2 v2.2.1 was used to build the index of the reference genome (GRCm39/mm39) and align clean paired end reads. Next, featureCounts v2.0.6 was used to count the read numbers mapped to each gene. The Fragments Per Kilobase of transcript sequenced per Million base pairs sequenced (FPKM) of each gene was calculated based on the length of the gene and read counts mapped to each gene. Gene expression analyses were completed with Novogene’s platform, Novomagic. R-squared values of log-transformed FPKMs between biological replicates were greater than 0.96 (Figures S6). Encode suggests that the Pearson correlation coefficient R^2^ should be greater than 0.92 (ENCODE Project Consortium, 2004). Differential expression analysis was performed using the DESeq2 R package v1.42.0. The resulting P values were FDR-adjusted using the Benjamini-Hochberg method. Genes with an adjusted P value (Q value) < 0.05 were considered differentially expressed. Gene Ontology (GO) enrichment analysis of differentially expressed genes (DEGs) was implemented by the clusterProfiler v4.8.1, in which gene length bias was corrected. All DEGs with Q < 0.05 were used for GO enrichment analysis. GO terms with Q values < 0.05 were considered significantly enriched by DEGs. Significant GO terms with the highest gene ratio were selected for plotting. DEGs were also submitted for Qiagen Ingenuity Pathway Analysis (IPA) using the Ingenuity Knowledge Base (Qiagen Bioinformatics). The fold change and Q values (< 0.05) of each DEG were used to perform core analysis, and IPA pathways with a Q value < 0.05 were considered significantly enriched. All mice used for bulk-sequencing studies were housed in Facility 2.

## Acknowledgements

We thank Dr. Samira Lawton and the team at TransnetYX for aid with the metagenomic sequencing of our murine fecal samples and for guidance on utilizing the OneCodex platform for data visualization. We thank the staff (especially Drs. Iris Herschmann and Vamseelatha Pakir) at the Stanford Human Immune Monitoring Center (HIMC), RRID:SCR_018266, for conducting Luminex plate assays to measure circulating serum cytokine abundance. We also thank Dr. Tyson Holmes (Statistical Director, HIMC) for initial consultation and insight on the use of the R Utility for batch and nonspecific binding correction for cytokine data analysis. We thank Dr. Joseph Garner for his statistical mentorship in Comparative Medicine 299 for further analysis of the serum cytokine data. All experimental paradigm figures were created in BioRender (Mehl, L. [2025]; https://BioRender.com). Finally, we thank the entirety of the Gibson and Giardino Labs past and present for their input on this project and specifically acknowledge Jacob Greene, Anna Badner, Sam H. Kim, Caroline Arellano-Garcia, Kelly Strickland, Allison Morningstar, Ivy Hoang, Brittany Bush, Ethan Rogers, Themis Tsarouchas, Jerry Cheng, Yohan Auguste, Daniela Rojo, Becca Buchanan, Sarah Wilson, and Tess Dierckx for their contributions to moving the project forward.

## Author contribution

L.C.M. and E.M.G. conceptualized the project. L.C.M., M.S.N., J.M.S., L.D.C., and A.G. executed various methodologies and collected data. L.C.M., M.S.N., J.M.S., Y.C., A.K., and E.M.G. analyzed and visualized the data. E.M.G. acquired funding. E.M.G. supervised the work. L.C.M. and E.M.G. wrote and edited the manuscript.

## Supplemental Figures

**Figure S1.**
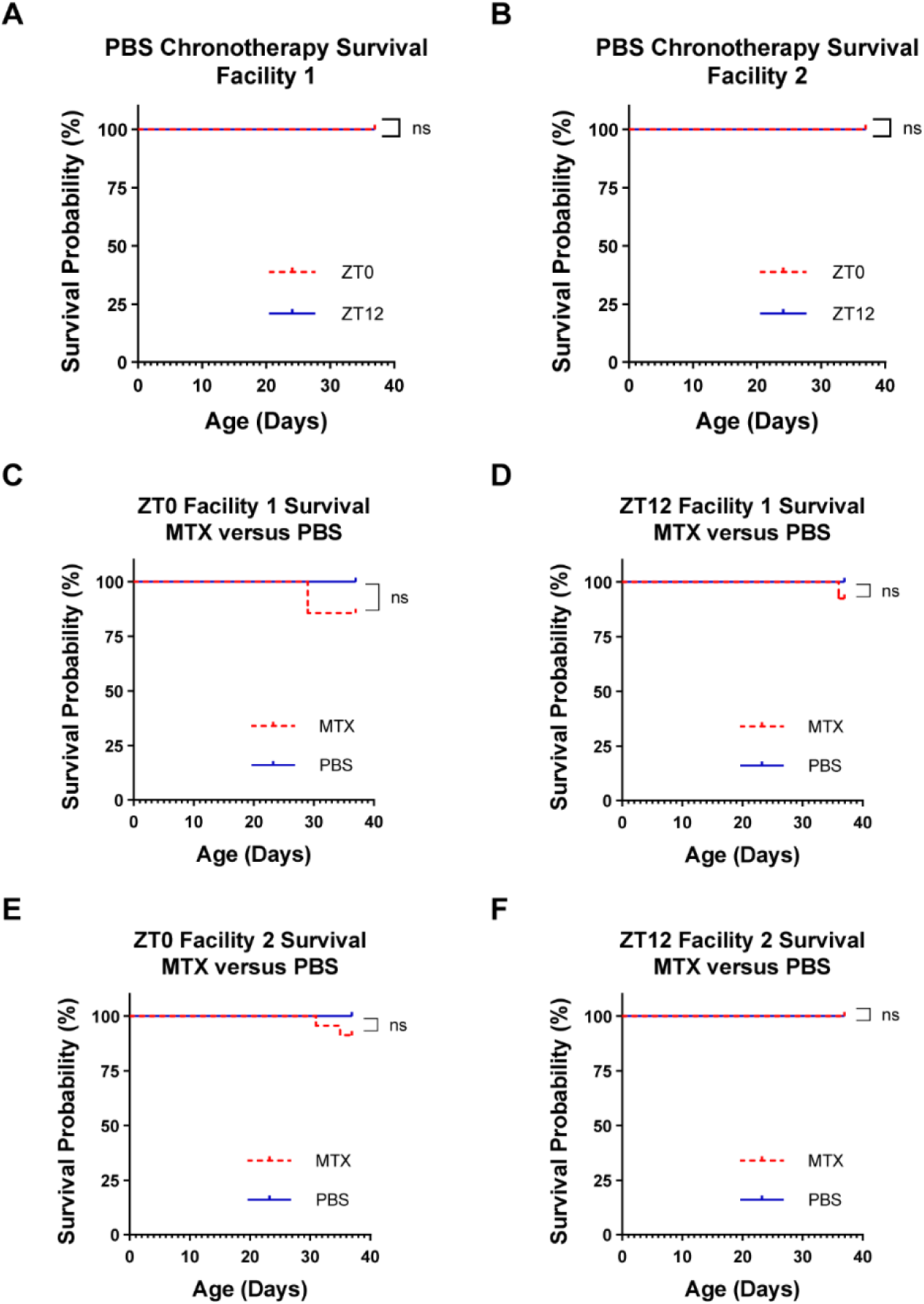
Chronotherapy does not adversely affect survival outcomes. (A) Animals treated with PBS in Facility 1 show no difference in overall survival after completion of the chronotherapy treatment paradigm when they are treated at ZT0 versus ZT12 (ZT0, n = 15; ZT12, n =13; p = >0.9999) by Log-rank (Mantel-Cox) test. (B) Animals treated with PBS in Facility 2 show no difference in overall survival after completion of the chronotherapy treatment paradigm when they are treated at ZT0 versus ZT12 (ZT0, n = 18; ZT12, n = 18; p = >0.9999) by Log-rank (Mantel-Cox) test. (C) Animals in Facility 1 treated at ZT0 show no difference in overall survival when they are treated with MTX versus PBS (PBS, n = 15; MTX, n = 14; p = 0.1360) by Log-rank (Mantel-Cox) test. (D) Animals in Facility 1 treated at ZT12 also show no difference in overall survival when they are treated with MTX versus PBS (PBS, n = 13; MTX, n = 13; p = 0.3173) by Log-rank (Mantel-Cox) test. (E) Animals in Facility 2 treated at ZT0 show no difference in overall survival when they are treated with MTX versus PBS (PBS, n = 18; MTX, n = 23; p = 0.2058) by Log-rank (Mantel-Cox) test. (F) Animals in Facility 2 treated at ZT12 also show no difference in overall survival when they are treated with MTX versus PBS (PBS, n = 18; MTX, n = 24; p = >0.9999) by Log-rank (Mantel-Cox) test.

**Figure S2.**
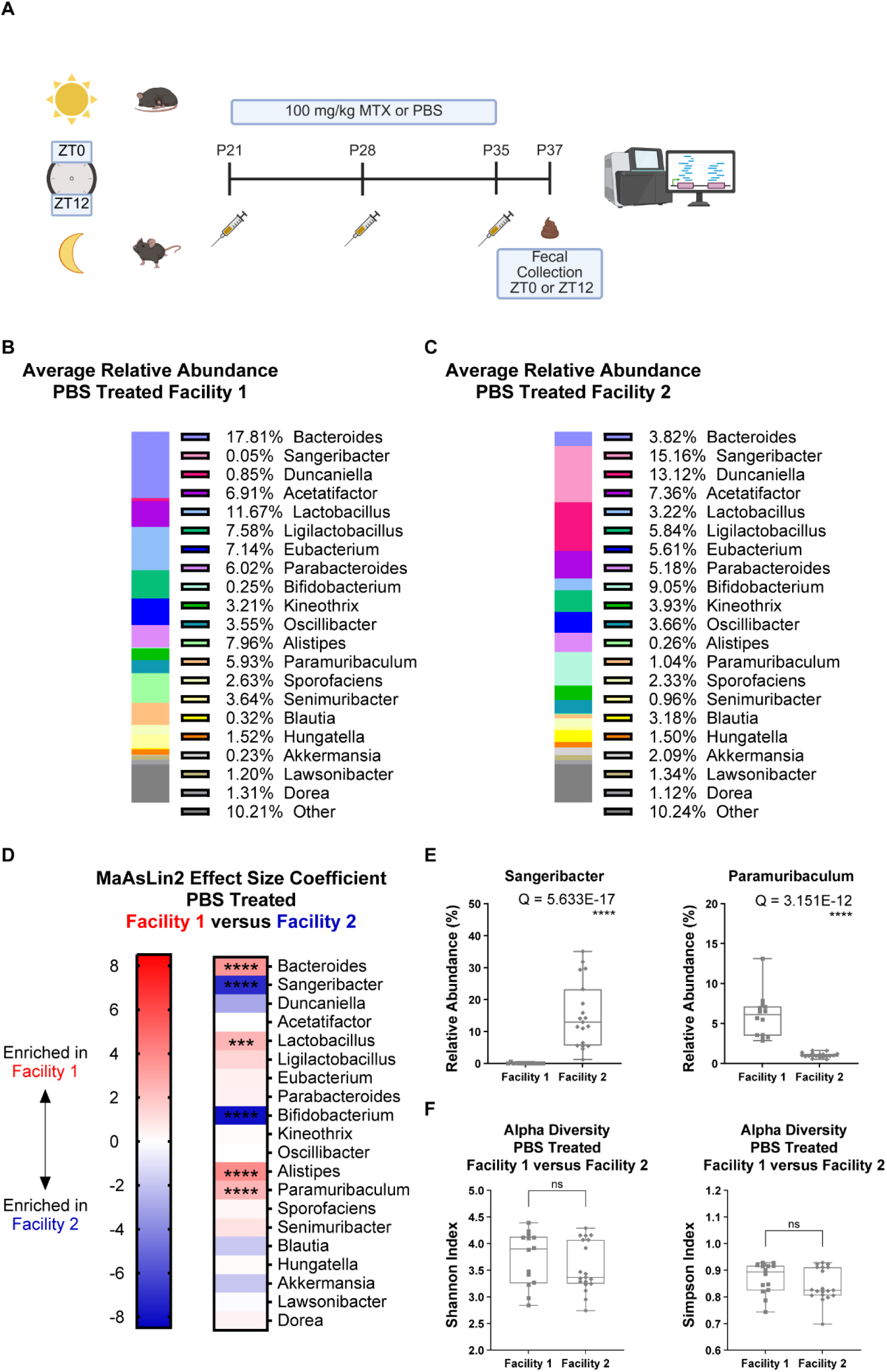
Housing facility determines microbiome composition at baseline. Animals were treated with PBS chronotherapy until P35 in either Facility 1 or Facility 2, and fecal pellets were collected at ZT0 and Zt12 on P37 for shotgun metagenomic microbiome sequencing. Sequencing reads were assigned to specific microbial genera and the relative abundance of microbes in each sample was computed using the OneCodex platform. The difference between microbial composition in Facility 1 versus Facility 2 after PBS treatment was assessed with the MaAsLin2 R package^60^ (See methods). (A) Microbiome sequencing paradigm - juvenile mice were treated at either ZT0 (lights on) or ZT12 (lights off) with either 100 mg/kg MTX or PBS via intraperitoneal injection starting at P21. They were treated once a week for 3 weeks followed by P37 fecal collection at the same time as injection. The feces were then sent to TransnetYX for shotgun metagenomic microbiome sequencing. Created in BioRender. Mehl, L. (2025) https://BioRender.com/fh0hfnn. (B) Average relative abundance of the top 20 microbial genera for PBS treated Facility 1 fecal samples (n = 14). (C) Average relative abundance of the top 20 microbial genera for PBS treated Facility 2 fecal samples (n = 19). (D) MaAsLin2 effect size coefficient was plotted as a heat map for each genus, with positive effect sizes (red) indicating enrichment in Facility 1 and negative effect sizes (blue) indicating enrichment in Facility 2 (Facility 1, n = 14; Facility 2, n = 19). Stars denote Q values of genera that were significantly affected by facility. A list of effect sizes and Q values for each genus can be found in Table S3. (E) Representative box and whisker plots of the relative abundance for genera significantly impacted by facility after PBS treatment as assessed by MaAsLin2 (Facility 1, n = 14; Facility 2, n = 19; Sangeribacter, Q = 5.633E-17; Paramuribaculum, Q = 3.151E-12). (F) The alpha diversity (both Shannon and Simpson indices) of each fecal sample was computed with the OneCodex Platform for samples from Facility 1 versus Facility 2 after PBS treatment. There was not a significant difference in alpha diversity when looking at either the Shannon Index (Facility 1, n = 14; Facility 2, n = 19; p = 0.3773) or the Simpson Index (Facility 1, n = 14; Facility 2, n = 19; p = 0.2120) by Mann-Whitney tests. n.s. p > 0.05; *p < 0.05; **p < 0.01; ***p < 0.001; ****p < 0.0001 n.s. Q > 0.05; *Q < 0.05; **Q < 0.01; ***Q < 0.001; ****Q < 0.0001

**Figure S3.**
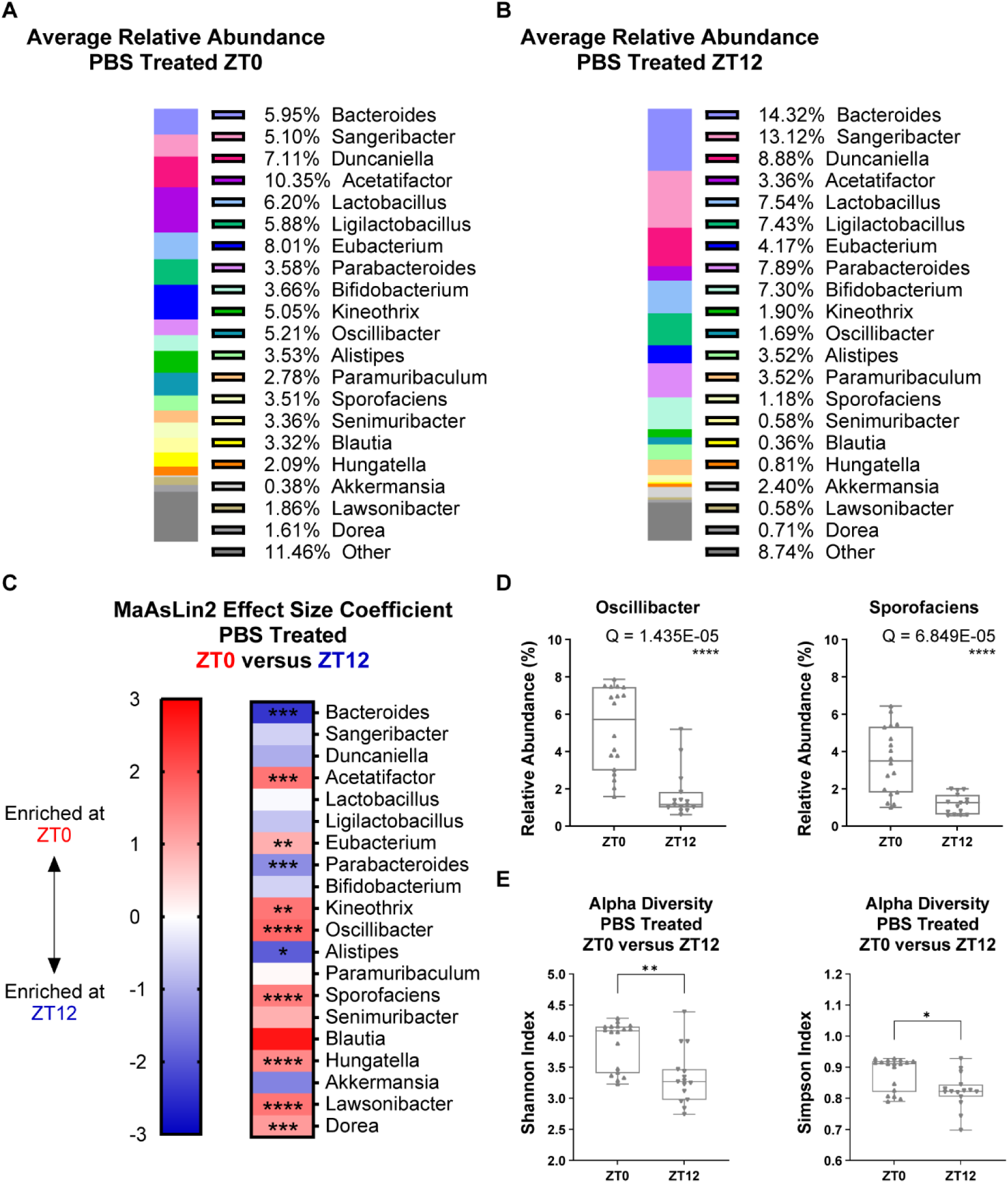
Time of day regulates microbiome composition at baseline. Animals were treated with PBS chronotherapy until P35 in Facility 1 and Facility 2, and fecal pellets were collected at ZT0 or ZT12 on P37 for shotgun metagenomic microbiome sequencing. Sequencing reads were assigned to specific microbial genera and the relative abundance of microbes in each sample was computed using the OneCodex platform. The difference between microbial composition at ZT0 versus ZT12 after PBS treatment was assessed with the MaAsLin2 R package^60^ (See methods). (A) Average relative abundance of the top 20 microbial genera for PBS treated ZT0 fecal samples (n = 18). (B) Average relative abundance of the top 20 microbial genera for PBS treated ZT12 fecal samples (n = 15). (C) MaAsLin2 effect size coefficient was plotted as a heat map for each genus, with positive effect sizes (red) indicating enrichment at ZT0 and negative effect sizes (blue) indicating enrichment at ZT12 (ZT0, n = 18; ZT12, n = 15). Stars denote Q values of genera that were significantly affected by time. A list of effect sizes and Q values for each genus can be found in Table S4. (D) Representative box and whisker plots of the Relative Abundance for genera significantly impacted by time after PBS treatment as assessed by MaAsLin2 (ZT0, n = 18; ZT12, n = 15; Oscillibacter, Q = 1.435E-05; Sporofaciens, Q = 6.849E-05). (E) The alpha diversity (both Shannon and Simpson indices) of each fecal sample was computed with the OneCodex Platform for samples at ZT0 versus ZT12 after PBS treatment. Alpha diversity was consistently higher at ZT0 versus ZT12 after PBS treatment when looking at both the Shannon Index (ZT0, n = 18; ZT12, n = 15; p = 0.0012) and the Simpson Index (ZT0, n = 18; ZT12, n = 15; p = 0.0132) by Mann-Whitney tests. n.s. p > 0.05; *p < 0.05; **p < 0.01; ***p < 0.001; ****p < 0.0001 n.s. Q > 0.05; *Q < 0.05; **Q < 0.01; ***Q < 0.001; ****Q < 0.0001

**Figure S4.**
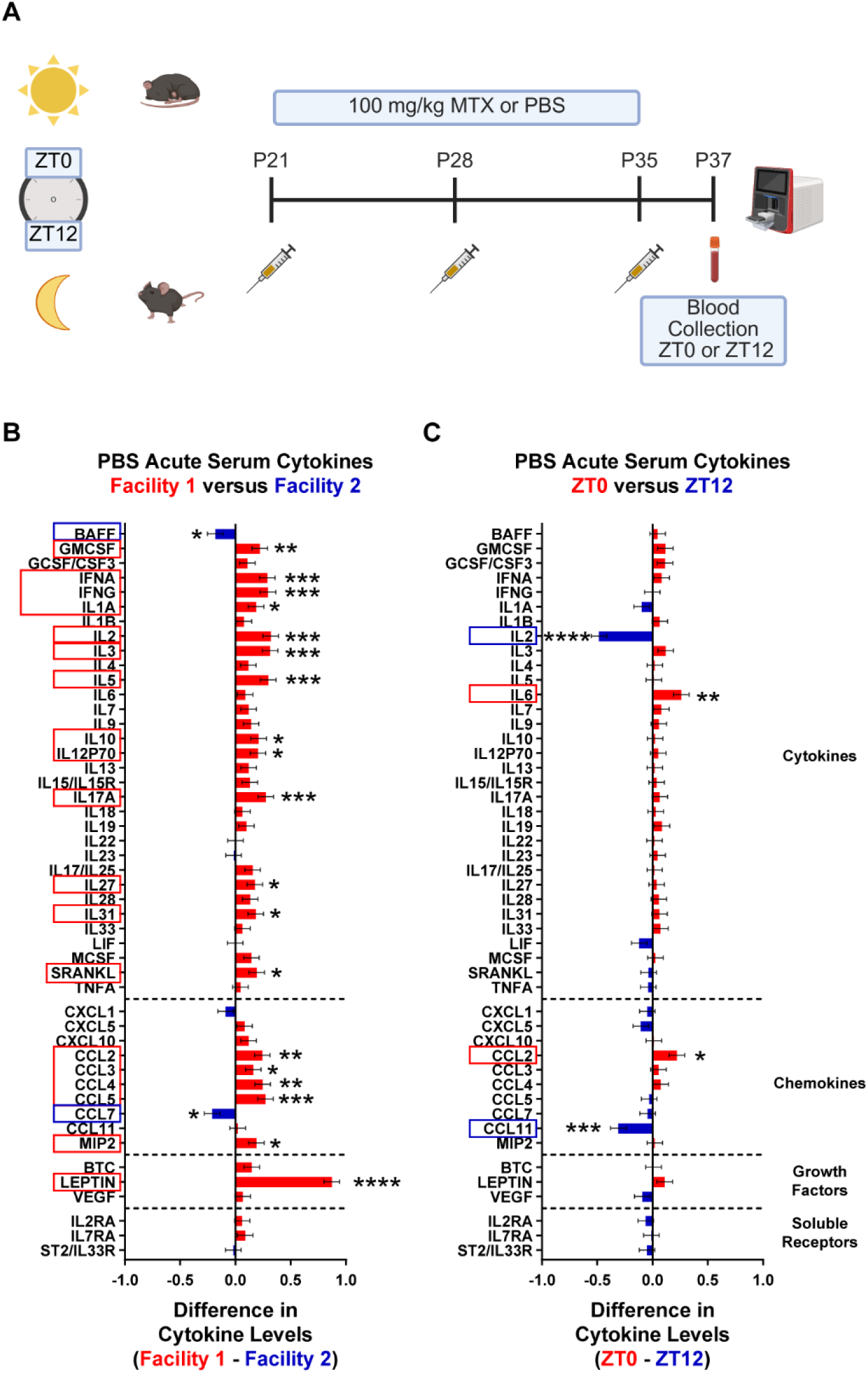
Housing facility and time alter acute peripheral inflammation at baseline. Animals were treated with PBS chronotherapy in Facility 1 or Facility 2, and blood was collected from the saphenous vein at either ZT0 or ZT12 on P37 at the same time as injection to assess peripheral inflammation. P37 PBS treated serum cytokine levels were analyzed via cytokine profiling analysis with an REML mixed effects model with Facility, Time, and Cytokine as fixed effects (See methods). (A) Acute P37 serum cytokine paradigm – In Facility 1 and Facility 2 juvenile mice were treated at either ZT0 (lights on) or ZT12 (lights off) with either 100 mg/kg MTX or PBS via intraperitoneal injection starting at P21. They were treated once a week for three weeks followed by P37 blood collection at the same time (ZT0 or ZT12) as injection. Blood was collected to measure circulating cytokine levels in the serum with a fluorescent bead-based assay. Created in BioRender. Mehl, L. (2025) https://BioRender.com/o7i8kno. (B) For PBS-treated samples collected at P37, a significant interaction between Facility and Cytokine was identified by cytokine profile analysis (Facility 1, n = 28; Facility 2, n = 23; p = < 0.0001; Table S5). P values for interactions between Facility and individual cytokines were FDR-adjusted, and cytokines with Q values less than 0.05 were considered significantly affected by interaction with Facility. Figure displays the difference in least squares mean estimates of Facility 1 versus Facility 2 (Facility 1, n = 28; Facility 2, n = 23). Values greater than zero indicate higher expression in Facility 1 (red), and values less than zero indicate higher expression in Facility 2 (blue). Data shown as ± SE. All cytokine results reported in Table S11. (C) For PBS-treated samples collected at P37, a significant interaction between Time and Cytokine was identified by the cytokine profile analysis (ZT0, n = 27; ZT12, n = 24; p = <0.0001; Table S5). P values for interactions between Time and individual cytokines were FDR-adjusted, and cytokines with Q values less than 0.05 were considered significantly affected by interaction with Time. Figure displays the difference in least squares mean estimates of ZT0 versus ZT12 (ZT0, n = 27; ZT12, n = 24; IL2, Q = 6.54E-10; CCL11, Q = 3.08E-4; IL6, Q = 0.00361; CCL2, Q = 0.0198). Values greater than zero indicate higher expression at ZT0 (red), and values less than zero indicate higher expression at ZT12 (blue). Data shown as ± SE. All cytokine results reported in Table S12.

**Figure S5.**
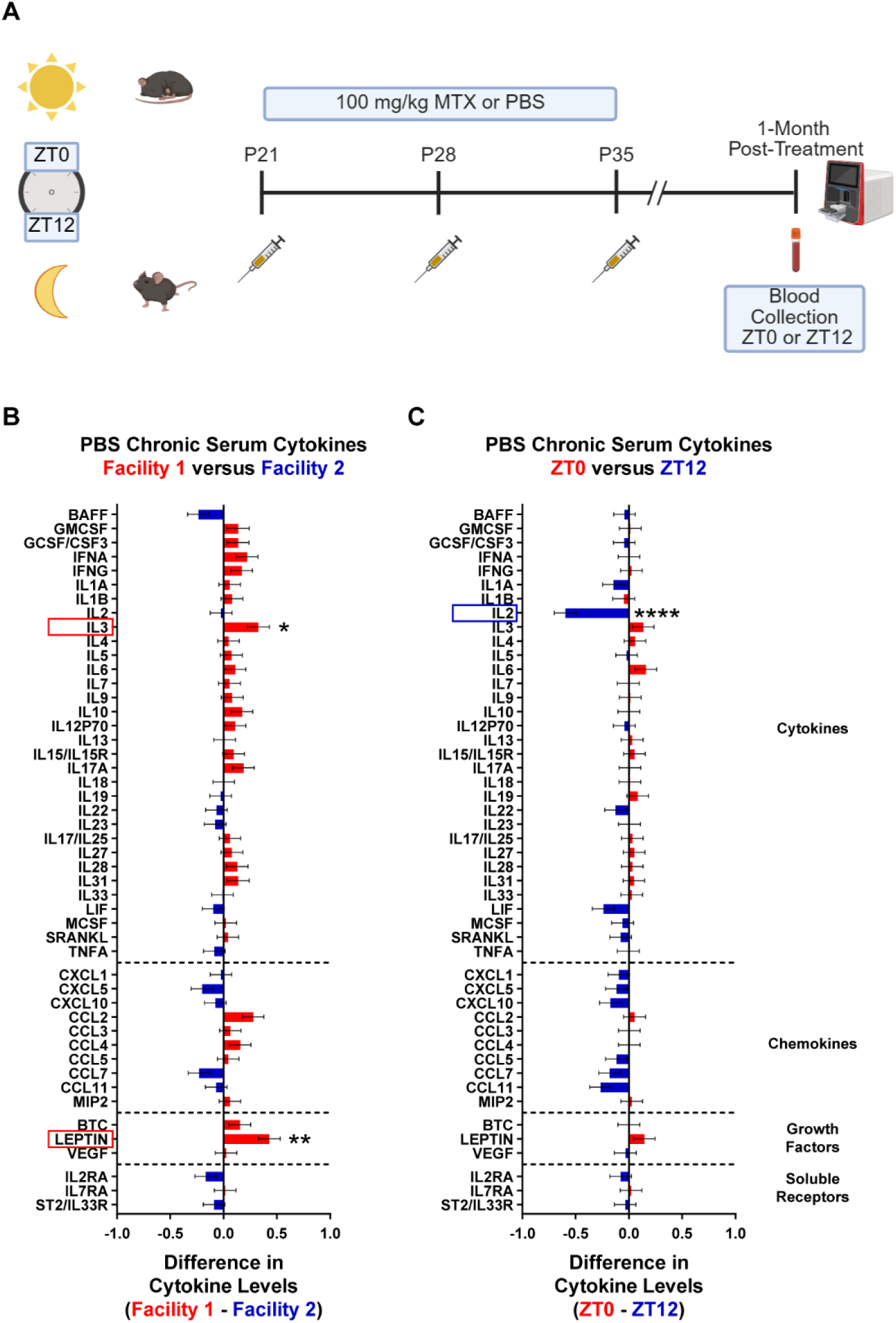
Housing facility and time have mild effects on chronic peripheral inflammation at baseline. Animals were treated with PBS chronotherapy in Facility 1 or Facility 2, and blood was collected from the saphenous vein at either ZT0 or ZT12 on P65 at the same time as injection to assess peripheral inflammation. P65 PBS treated serum cytokine levels were analyzed via cytokine profiling analysis with an REML mixed effects model with Facility, Time, and Cytokine as fixed effects (See methods). (A) Chronic P65 serum cytokine paradigm – In Facility 1 and Facility 2 juvenile mice were treated at either ZT0 or ZT12 with either 100 mg/kg MTX or PBS via intraperitoneal injection starting at P21. They were treated once a week for three weeks followed by P65 blood collection at the same time (ZT0 or ZT12) as injection. Blood was collected to measure circulating cytokine levels in the serum with a fluorescent bead-based assay. Created in BioRender. Mehl, L. (2025) https://BioRender.com/v6el5e9. (B) For PBS-treated samples collected at P65, a significant interaction between Facility and Cytokine was identified by the cytokine profile analysis (Facility 1, n = 22; Facility 2, n = 15; p = <0.0001; Table S5). P values for interactions between Facility and individual cytokines were FDR-adjusted, and cytokines with Q values less than 0.05 were considered significantly affected by interaction with Facility. Figure displays the difference in least squares mean estimates of Facility 1 versus Facility 2 (Facility 1, n = 22; Facility 2, n = 15); Leptin, Q = 0.00141; IL3, Q = 0.0336). Values greater than zero indicate higher expression in Facility 1 (red), and values less than zero indicate higher expression in Facility 2 (blue). Data shown as ± SE. All cytokine results reported in Table S14. (C) For PBS-treated samples collected at P65, a significant interaction between Time and Cytokine was identified by the cytokine profile analysis (ZT0, n = 20; ZT12, n = 17; p = 0.0039; Table S5). P values for interactions between Time and individual cytokines were FDR-adjusted, and cytokines with Q values less than 0.05 were considered significantly affected by interaction with Time. Figure displays the difference in least squares mean estimates of ZT0 versus ZT12 (ZT0, n = 20; ZT12, n = 17; IL2, Q = 3.25E-7). Values greater than zero indicate higher expression at ZT0 (red), and values less than zero indicate higher expression a ZT12 (blue). Data shown as ± SE. All cytokine results reported in Table S15. n.s. Q > 0.05; *Q < 0.05; **Q < 0.01; ***Q < 0.001; ****Q < 0.0001

**Figure S6.**
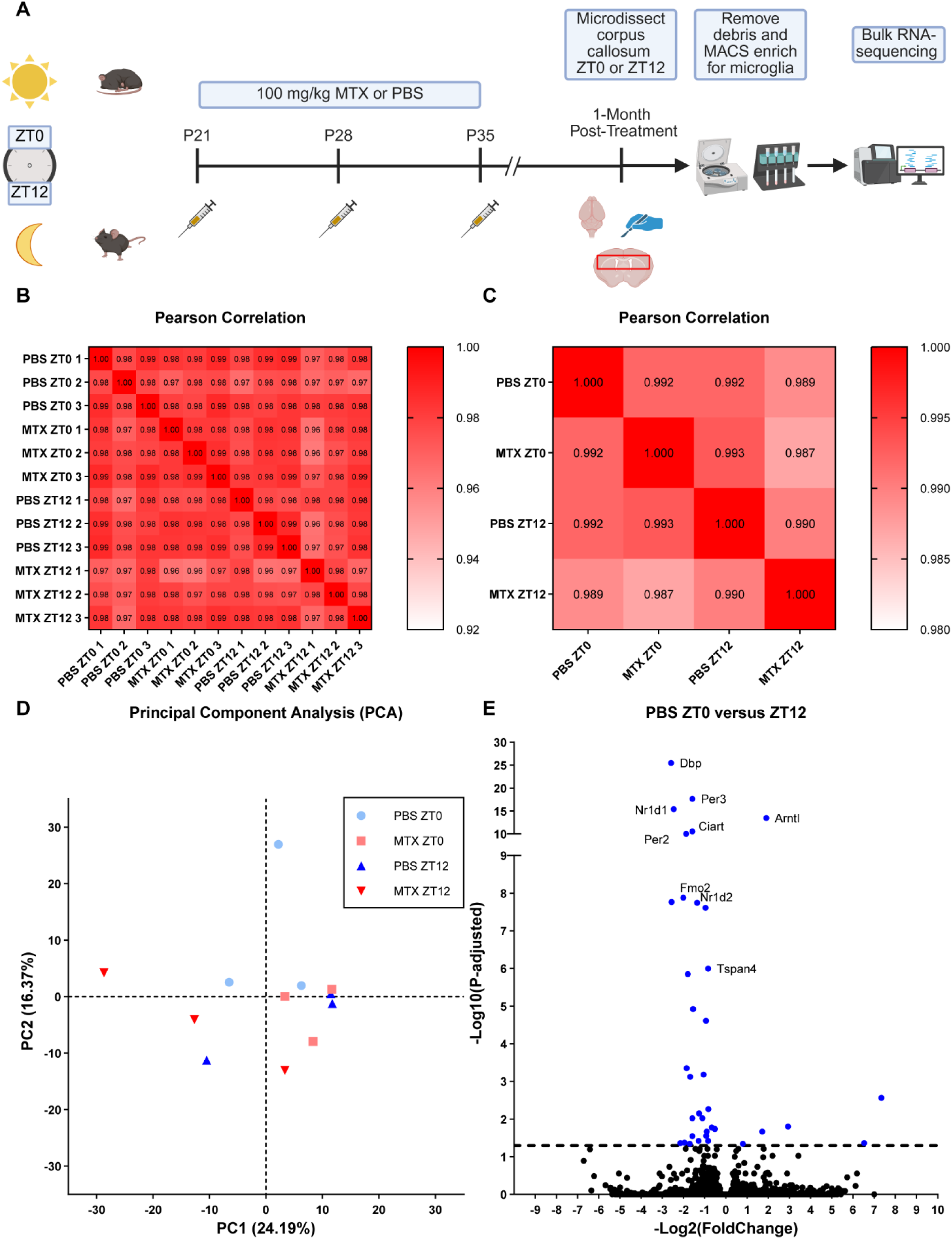
Bulk RNA-sequencing of white matter microglia after chronotherapy. Bulk RNA-sequencing (RNA-seq) was performed at 1-month post-treatment on MACS enriched microglia from the corpus callosum of mice that underwent the chronotherapy treatment paradigm in Facility 2. (A) Chronotherapy RNA-sequencing paradigm - juvenile mice were treated at either ZT0 (lights on) or ZT12 (lights off) with either 100 mg/kg MTX or PBS once a week for three weeks. At 1-month post-treatment, at the same time as injection (ZT0 or ZT12), the corpus callosum was microdissected, and microglia were isolated via MACS for bulk RNA-sequencing. Created in BioRender. Mehl, L. (2025) https://BioRender.com/hlltubx. (B) Pearson correlation coefficient matrix with biological replicate distances. (C) Pearson correlation coefficient matrix with biological group distances (n = 3 biological replicates per biological group). (D) Principal component analysis (PCA) on the gene expression values (FPKM) for each biological group (n = 3 biological replicates per biological group). (E) Volcano plot showing differential expressed genes in PBS ZT0 versus ZT12 samples. Genes with P-adjusted (Q value) less than 0.05 are shown in dark blue (n = 3 biological replicates per biological group).

